# *rbfox1* LoF mutants show disrupted *bdnf/trkb2* and *crhb/nr3c2* expression and increased cortisol levels during development coupled with signs of allostatic overload in adulthood

**DOI:** 10.1101/2024.10.09.616976

**Authors:** Adele Leggieri, Judit García-González, Saeedeh Hosseinian, Peter Ashdown, Sofia Anagianni, Xian Wang, William Havelange, Noèlia Fernàndez-Castillo, Bru Cormand, Caroline H. Brennan

**Affiliations:** Centre for Brain and Behaviour, School of Biological and Behavioural Sciences, Queen Mary University of London, Mile End Rd, London, E1 4NS, United Kingdom; Department of Genetics and Genomic Sciences, Icahn School of Medicine, Mount Sinai, New York City, NY 10029, USA; Departament de Genètica, Microbiologia i Estadística, Facultat de Biologia, Universitat de Barcelona, Barcelona, Catalunya, 08028, Spain; Centro de Investigación Biomédica en Red de Enfermedades raras (CIBERER), Spain; Institut de Biomedicina de la Universitat de Barcelona, Barcelona, Catalunya, 08028, Spain; Institut de recerca Sant Joan de Déu, Espluges de Llobregat, Catalunya, 08950, Spain

**Author notes:** **Correspondence:** Caroline H. Brennan.

## Abstract

Mutations in the *RBFOX1* gene are associated with psychiatric disorders but how RBFOX1 influences psychiatric disorder vulnerability remains unclear. Recent studies showed that RBFOX proteins mediate the alternative splicing of PAC1, a critical HPA axis activator. Further, RBFOX1 dysfunction is linked to dysregulation of BDNF/TRKB, a pathway promoting neuroplasticity, neuronal survival, and stress resilience. Hence, RBFOX1 dysfunction may increase psychiatric disorder vulnerability via HPA axis dysregulation, leading to disrupted development and allostatic overload. To test this hypothesis, we generated a zebrafish *rbfox1* loss of function (LoF) line and examined behavioural and molecular effects during development. We found that *rbfox1* LoF mutants exhibited hyperactivity, impulsivity and heightened arousal, alongside alterations in proliferation – traits associated with neurodevelopmental and stress-related disorders. In adults, loss of *rbfox1* function led to decreased fertility and survival, consistent with allostatic overload. At the molecular level, at larval stages *rbfox1* mutants showed increased cortisol levels and disrupted expression of key stress-related genes (*bdnf, trkb2, pac1a-hop, crhb, nr3c2*). Pharmacological intervention targeting TRKB restored *crhb* and *nr3c2* gene expression and hyperactive and hyperarousal behaviours. In adults, dysregulation of *crhb, nr3c2* and *bdnf/trkb2* genes was only seen following acute stress exposure. Our findings reveal a fundamental role for RBFOX1 in integrating stress responses through its regulation of BDNF/TRKB and neuroendocrine signalling.

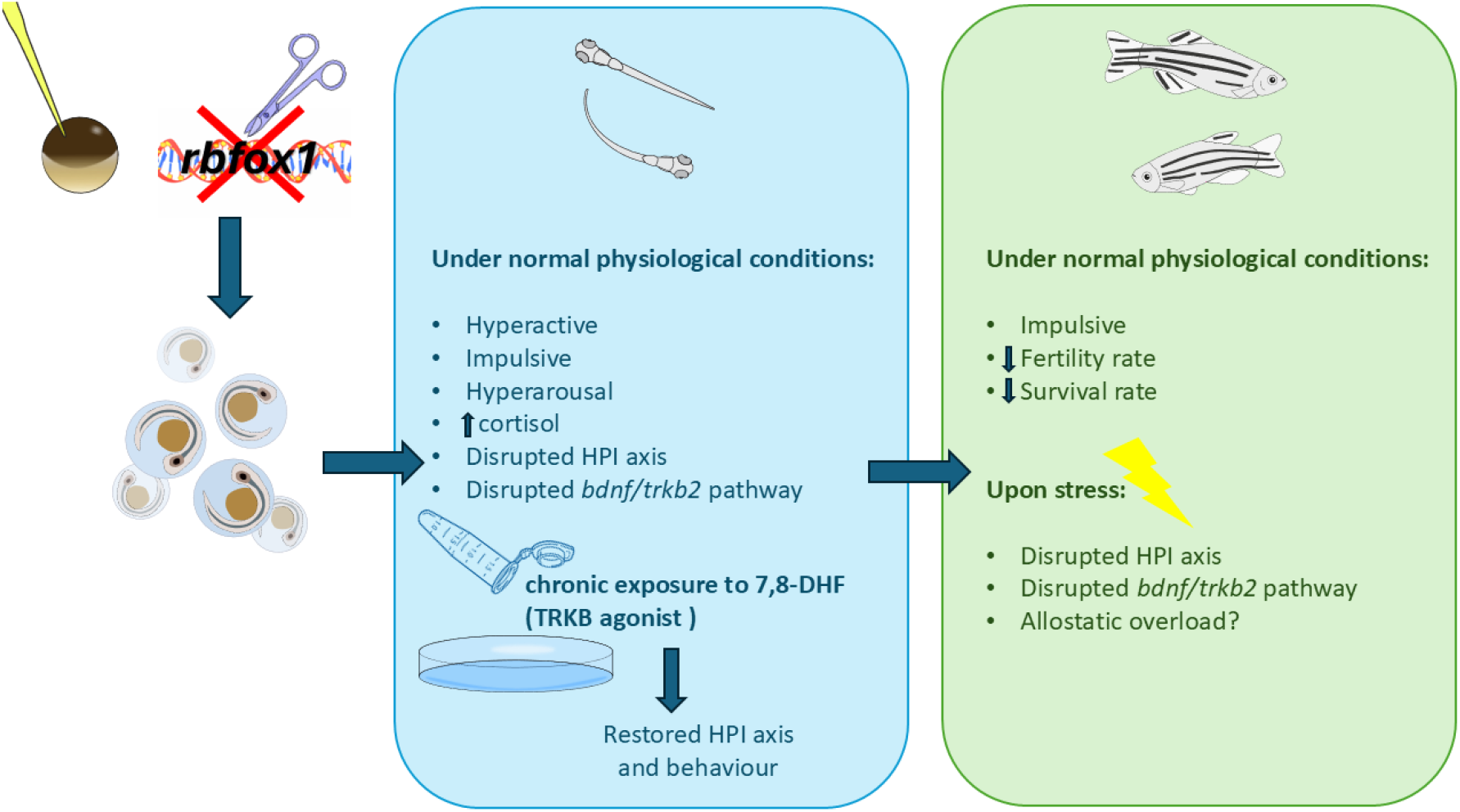

## INTRODUCTION

Psychiatric disorders are influenced by multiple genes, with a substantial portion of their heritability linked to common genetic variations^1^. Identifying gene variants associated with multiple psychiatric disorders is crucial for understanding their shared genetic underpinnings and enhancing therapeutic strategies.

RNA binding fox-1 homologue 1 (RBFOX1), also known as FOX1 or ataxin 2-binding protein 1 (A2BP1), is a splicing factor highly conserved among vertebrates and expressed in the heart, brain and skeletal muscle, where it contributes to normal development and function^2,3^. Alterations in *RBFOX1* expression or function have been associated with susceptibility to psychiatric disorders, particularly those associated with changes in stress-related behaviour^4–6^. In a cross-disorder genome-wide association study (GWAS), *RBFOX1* emerged as the second most pleiotropic *locus*, showing association with seven out of the eight disorders studied: schizophrenia, bipolar disorder, depression, attention-deficit/hyperactivity disorder (ADHD), autism spectrum disorder (ASD), obsessive-compulsive disorder and Tourette syndrome^1^. In mice and zebrafish, *Rbfox1/rbfox1* knockout caused hyperactivity, increased anxiety-like behaviour and altered social behaviour^2,6^. However, the mechanisms by which *RBFOX1* genetic variants contribute to psychiatric disease are poorly understood.

In both mice and zebrafish, it has been suggested that Rbfox1 regulates the Pituitary Adenylate Cyclase-Activating Polypeptide Type 1 Receptor (Pac1) alternative splicing, an important mediator of corticotropin releasing hormone (*CRH*) synthesis in the hypothalamus^4,7^. *CRH* is the first hormone to be secreted in response to stress by the hypothalamic-pituitary-adrenal (HPA) axis (hypothalamic-pituitary-interrenal (HPI) axis in fish)^8^. In mammals there are several PAC1 isoforms with the predominant isoforms in the brain being PAC1-hop (long isoform) and PAC1-short^9^. In mice and zebrafish, following acute stress exposure, both *Pac1-hop/pac1-hop* and *Pac1-short/pac1-short* expression increased, while at the late recovery phase, only Pac1-hop was still up-regulated^4^.

Another regulator of the stress response and a resilience factor against chronic stress-induced psychopathology is the brain derived neurotrophic factor (BDNF)/Tropomyosin receptor kinase B (TRKB) pathway^10–12^. BDNF is a neurotrophin possessing a pivotal role in the modulation of neurotransmission and synaptic plasticity and dysregulation of the BDNF/TRKB pathway has been associated with several neuropsychiatric diseases, including anxiety/stress disorders^13,14^. In human neural stem cells, *RBFOX1* knockdown increased *BDNF* expression levels^15^, while another study in mice identified *TrkB* as a target of RBFOX1 within the hippocampus^16^. Further, PAC1-short activation can elevate *BDNF* levels and potentiate TRKB activity, enhancing BDNF/TRKB neuroprotective and plasticity-promoting effects, especially in the context of stress response and neuropsychiatric health^17^.

Given that both PAC1 and BDNF/TRKB influence how the brain adapts to and manages stress, RBFOX1 variants may increase susceptibility to psychiatric disorders through dysregulation of the stress response, leading to adaptive plasticity and disrupted development in the short-term and allostatic overload in the long-term. Allostasis refers to the collective processes by which the body actively maintains stability (or homeostasis) in response to environmental stressors^12^. When allostasis primary mediators (e.g., HPA axis hormones, catecholamines, cytokines) are overactivated or fail to return to normal, it leads to an allostatic state^18,19^. The cumulative results of an allostatic state are referred to as allostatic load and, when maladaptive, allostatic overload^18^. Excess glucocorticoid exposure during early life and early life stress have been shown to cause prolonged activation of the allostatic systems, ultimately leading to allostatic overload^19–22^, while neurotrophic factors such as BDNF play a key role in regulating adaptive plasticity and mechanisms counteracting damage caused by allostatic overload^12^. Here, to explore the possibility that *RBFOX1* loss of function (LoF) leads to increased vulnerability to psychiatric disease through HPA axis hyperactivation and allostasis-induced adaptation during development, we generated a zebrafish line carrying a predicted *rbfox1* LoF mutation using CRISPR-Cas9 gene editing and assessed behavioural and molecular changes at different developmental stages. We hypothesized that RBFOX1 regulates HPA axis activity through an effect on BDNF/TRKB signalling leading to disrupted brain development.

## RESULTS

### Generation of a loss of function rbfox1 line

CRISPR-Cas9 genome editing generated a 19 base pair deletion (NM_001005596.1, nucleotides 120-138, TCCCATCGGCCCAGTTCGC) that introduced a premature termination codon (PTC) at position 58 in the *rbfox1* amino acid sequence. This line is recorded on ZFIN as qm4, ZDB-ALT-240222-6. Details regarding the generation of the line, nucleotide and amino acid sequences can be found in our previous study^2^. Wild type animals are denoted throughout as *rbfox1*^*+/+*^, heterozygous mutants as *rbfox1*^*+/19del*^ and homozygous mutants as *rbfox1*^*19del/19del*^.

To evaluate whether the PTC elicited mRNA non-sense mediated decay (NMD) and consequent reduction of *rbfox1* mRNA in mutant fish, we examined *rbfox1* expression by quantitative Real-Time polymerase chain reaction (qPCR) and by *in situ* hybridisation (ISH). As *RBFOX1* itself is alternatively spliced to generate nuclear and cytoplasmic isoforms^7^, we designed primers targeting all *rbfox1* zebrafish isoforms available on NCBI (see Supplementary Table1). qPCR showed that rbfox1 transcript levels were significantly lower in mutant larvae compared to *rbfox1*^*+/+*^ siblings (p_*rbfox1*_ *+/+* _*vsrbfox1*_*+/19del* < 0.05, p _*rbfox1*_*+/+* _vs rbfox1_ ^19*del*/19*del*^ < 0.01), and ISH showed that *rbfox1* was not detectable in rbfox1^19del/19del^ fish, at either larval or adult stages (Supplementary Figure 1A-C).

These results confirm degradation of defective *rbfox1* mRNA in mutant fish.

### rbfox1 mutant fish show hyperactivity, impulsivity and hyperarousal behaviour

As mutations in *RBFOX1* locus have been linked to several psychiatric diseases in humans, we assessed zebrafish larvae and adult fish for phenotypic traits associated with such disorders and examined rbfox1 mRNA distribution in *rbfox1*^*+/+*^ larvae and adults (Tübingen strain).

In humans, RBFOX1 copy number variants (CNVs) and LoF mutations are risk factors for ADHD6. Two major ADHD traits are hyperactivity and increased impulsivity. We therefore assessed hyperactive and impulsive behaviour of *rbfox1*^*+/+*^, *rbfox1*^*+/19del*^ and *rbfox1*^*19del/19del*^ 5 days post fertilization (dpf) larvae and adults. When we measured larval locomotion, we observed a gene dosage effect on distance travelled, whereby rbfox1 mutant larvae travelled greater distances than wild type siblings (p _*rbfox1*_*+/+* _vs rbfox1_^*+/19del*^ < 0.05, p _*rbfox1*_*+/+* _vs rbfox1_^*19del /19del*^ < 0.0001, p _*rbfox1*_^*+/19del*^ _vs rbfox1_^*19del /19del*^ < 0.05) (Supplementary Figure 2A), and a significant increase in the swimming speed of rbfox1^*19del/19del*^ larvae (p _*rbfox1*_ *+/+* _vs rbfox1_ ^*19del/19del*^ < 0.0001, p _*rbfox1*_ ^*+/19del*^ _vs rbfox1_ ^*19del/19del*^ < 0.05) (Supplementary Figure 2B). This is in line with hyperactivity observed in adult rbfox1^*19del/19del*^, in our previous study and in other *rbfox1* model2,23. We also measured larval burst swimming, a parameter previously used as a measure to predict impulsive behaviour in zebrafish larvae24. We found a significant increase in the number of peaks (acceleration events when the fish travelled > 5 mm in < 12 sec) in *rbfox1*^*19del/19del*^ larvae (p _*rbfox1*_*+/+* _vs rbfox1_^*19del /19del*^ < 0.0001, p _*rbfox1*_^*+/19del*^ _vs rbfox1_^*19del /19del*^ < 0.0001) (Figure 1A). Impulsive behaviour was then assessed in adult (7 months old) fish using the 5-choice serial reaction time task (5-CSRTT)25. The 5-CSRTT assay measures sustained attention and impulsive action by requiring an animal to detect a brief visual cue (white light stimulus) presented randomly across one of five apertures and “nosepoke” that aperture to signal recognition (Supplementary Figure 3). If correct, a “food” signal light comes on (at the opposite end of the test chamber) and the animal collects food reward (Supplementary Figure 3). Crucially, a pause occurs prior to the onset of the stimulus light (called a pre-stimulus interval [PSI]), during which a ‘premature’ response can be interpreted as ‘impulsivity’-impulsive action25. The assay consists of five stages (Supplementary Table2) each run for at least a week until fish are promoted to the next stage. We found that 79% *rbfox1*^*+/+*^, 72% *rbfox1*^*+/19del*^ and 62% *rbfox1*^*19del/19del*^ learned the task within 9 weeks. We found no significant differences in the correct responses (stages 2-5, Figure 1B-E). In stage 5, we found a significant difference in the number of premature responses (increased impulsivity) such that *rbfox1*^*19del/19del*^ fish were more impulsive compared to rbfox1*+/+* siblings (p < 0.05) (Figure 1F).

**Figure 1.**
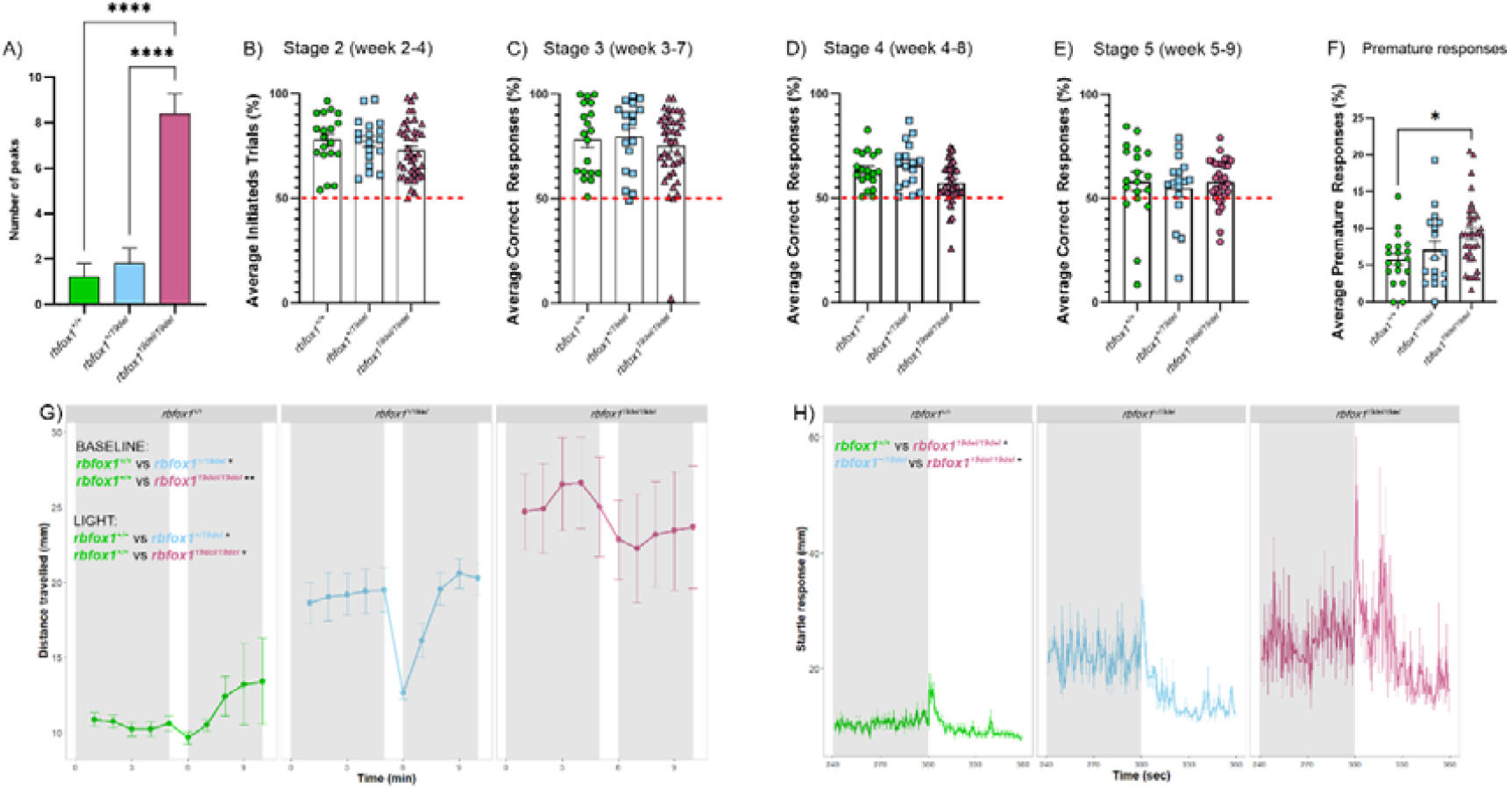
*rbfox1* mutant fish show hyperactivity, impulsivity and hyperarousal behaviour. **A** number of peaks/swimming burst (acceleration events when the fish travelled > 5 mm in < 12 s); N = *rbfox1+/+* 24; *rbfox1*^*+/19del*^ 24; *rbfox1*^*19del/19del*^ 24. **B-F**) 5-choice serial reaction time task in adult (7 months old) zebrafish (rbfox1^+/+^, rbfox1^*+/19del*^, *rbfox1*^*19del/19del*^): **B**) number of initiated trials during stage 2 (weeks 2-4); all *rbfox1+/+* (19), all *rbfox1*^*+/19del*^ (18) and all *rbfox1*^19del/19del^ (40) moved to the next stage; **C-E)** average correct responses for stages 3-5 (weeks 3-9); **C)** at the end of stage 3 (weeks 3-7), all rbfox1*+/+* (19), 17 *rbfox1*^*+/19del*^ and 39 *rbfox1*^*19del/19del*^ moved to the next stage; **D)** at the end of stage 4 (weeks 4-8), all rbfox1*+/+* (19), all *rbfox1*^*+/19del*^ (17) and 33 *rbfox1*^*19del/19del*^ moved to the next stage; **E)** at the end of stage 5 (weeks 5-9), 15 *rbfox1+/+*, 13 *rbfox1*^*+/19del*^ and all *rbfox1*^*19del/19del*^ (33) showed ≥ 50% correct responses; **F)** premature responses during stage 5 (weeks 5-9). Each dot, square or triangle in D-H) represents a single adult zebrafish *rbfox1+/+, rbfox1*^*+/19del*^ and *rbfox1*^*19del/19del*^ respectively; N = *rbfox1+/+* 19; *rbfox1+/19del* 18; *rbfox1*^*19del/19del*^ 40. **G-H)** Forced light-dark transition assay in 5dpf zebrafish larvae (^rbfox1+/+^, rbfox1^*+/19del*^, *rbfox1*^*19del/19del*^): **G)** distance travelled during the 10 min assay; **H)** 1 sec time bin resolution plots of the dark-light transition; N = *rbfox1+/+* 43; *rbfox1+/19del* 34; *rbfox1*^*19del/19del*^ 37. All larvae employed in behavioural experiments were progeny of *rbfox1*^*+/19del*^ in-cross and were genotyped after experiments and prior to data analysis. In all graphs: bars represent standard error of the mean (SEM); * p < 0.05; ** p < 0.01; *** p < 0.001; **** p < 0.0001.

*RBFOX1* CNVs have also been identified in individuals with schizophrenia6,15. Deficits in habituation to acoustic startle reflex are seen in both schizophrenic patients and animal models of schizophrenia26. The acoustic startle assay has been widely employed to measure habituation in animals, including zebrafish27. The magnitude of the startle response and the extent of habituation can also serve as indicators of hyperarousal linked to heightened stress responses28. We therefore tested 5dpf zebrafish larvae (*rbfox1*^*+/+*^, *rbfox1*^*+/19del*^ and *rbfox1*^*19del/19del*^) in the habituation to acoustic startle response assay. Consistent with our hyperactivity assay, during the baseline (first 10 min), the genotype had a significant main effect on distance travelled [Effect of genotype: χ2(2) = 8.3397, p < 0.05] whereby *rbfox1*^*19del/19del*^ larvae travelled greater distances than *rbfox1*^*+/+*^ siblings (p < 0.05). During the startle stimuli, we observed a main effect of genotype [Effect of genotype: χ2(2) =33.536, p < 0.0001] and stimulus number [Effect of stimulus number: χ2(9) = 487.968, p < 0.0001] on distance travelled, and a significant two-way interaction between genotype and stimulus number [Effect of genotype by stimuli: χ2(18) = 31.511, p < 0.05], whereby *rbfox1*^*19del/19del*^ larvae travelled greater distances than *rbfox1*^*+/+*^ (p < 0.001) and rbfox1*+/19del* (p < 0.01) (Supplementary Figure 2C). When assessed for the rate of habituation, wild type larvae showed a habituation response to repeated acoustic startle consistent with previous reports29: 100% of *rbfox1*^*+/+*^animals responded to the first acoustic stimulus, but only 22% responded to the last. When we examined the rate of habituation over time, in line with previous findings30, we observed a significant genotype effect [Effect of genotype: χ2(2) =7.2676, p < 0.05] and a significant two-way interaction between genotype and stimulus number [Effect of genotype by stimulus number: χ2(18) =132.8476, p < 0.001] whereby *rbfox1*^*19del/19del*^ showed reduced rate of habituation and a greater proportion of responders compared to *rbfox1*^*+/+*^ siblings (p _*rbfox1*_*+/+* _vs rbfox1_^*19del/19del*^ < 0.05) (Supplementary Figure 2D).

SNPs in *RBFOX1* locus have also been associated with anxiety6,31. Hence, we assessed anxiety-like behavior in 5dpf zebrafish larvae using the forced light-dark transition (FLDT) assay and in adult zebrafish using the novel tank diving (NTD) assay. In the FLDT assay, zebrafish are exposed to sudden transitions in illumination with effects on locomotion and amplitude of response on transition from dark to light being used as a measure of anxiety-like behaviour: the increased locomotion/startle response upon the light-dark transition is a measure of anxiety-like behaviour29,32,33. Consistent with our acoustic startle assay, we found that during the baseline (first 5 min) (Figure 1G) genotype had a significant main effect on distance travelled [Effect of genotype: χ2(2) = 14.4878, p < 0.001] whereby *rbfox1*^*19del*^ larvae travelled greater distances than *rbfox1*^*+/+*^ siblings (*p* _*rbfox1*_^*+/+*^ _vs rbfox1_^*+/19del*^ < 0.05, p _*rbfox1*_^*+/+*^ _*vs rbfox1*_^*19del/19del*^ < 0.01). During the 1min light flash (min 5-6 of the assay) we found a significant main effect of genotype on distance travelled [Effect of genotype: χ2(2) = 8.8518, p < 0.05] with *rbfox1*^*19del*^ larvae travelling greater distances than *rbfox1*^*+/+*^ siblings (p _*rbfox1*_^*+/+*^ _*vs rbfox1*_^*+/19del*^ < 0.05, p _*rbfox1*_^*+/+*^ _*vs rbfox1*_^*19del/19del*^ < 0.05) (Figure 1H). On transition from dark to light (analysed at 1 sec time bin resolution from sec 240 to sec 360 as the startle response during the transition is visible only at this resolution) we found a significant main effect of time [Effect of time: χ2(118) = 1051.3482, p < 0.001] and genotype [Effect of genotype: χ2(2) = 8.9155, p < 0.05], and a significant two-way interaction between time and genotype [Effect of time by genotype: χ2(236) = 514.8308, p < 0.001] on the amplitude of response, whereby *rbfox1*^*19del/19del*^ larvae startled more than *rbfox1*^*+/+*^ (p < 0.05) and *rbfox1*^*+/19del*^ (p < 0.05) siblings (Figure 1H). As there was a significant difference in basal locomotion, we also examined the amplitude of response on transition from dark to light normalizing the data against baseline, as reported previously34. We observed similar results as in absence of normalisation: significant main effect of time [Effect of time: χ2(118) = 1249.918, p < 0.001] and genotype [Effect of genotype: χ2(2) = 37.716, p < 0.001], and a significant two-way interaction between time and genotype [Effect of time by genotype: χ2(236) = 4532.966, p < 0.001] whereby *rbfox1*^*19del/19del*^ larvae startled more than *rbfox1*^*+/+*^ (p < 0.0001) and *rbfox1*^*+/19del*^ (p < 0.001) siblings.

Anxiety-like behaviour in adult animals was assessed using the NTD assay. When introduced to a novel tank, zebrafish will first dive to the bottom of the tank, to seek protection, and then gradually increase their swimming over time35. Over the entire duration of the NTD assay, we found no significant differences between rbfox1 genotypes(p > 0.05). However, during the first minute of the assay, we found a significant two-way interaction between genotype and the time spent at the bottom of the tank [Effect of genotype by proportion at the bottom tank: χ2(10) =22.3333, p < 0.05], such that *rbfox1*^*19del/19del*^ fish spent less time in the bottom than *rbfox1*^*+/+*^ fish (p < 0.001) (Supplementary Figure 2E). In line with previous findings2, we observed no significant differences in distance travelled (p > 0.05) (Supplementary Figure 3F), nor in the number of the transitions to the top area of the tank between rbfox1 genotypes (p > 0.05) (Supplementary Figure 3E).

Thus, our data showed that loss of rbfox1 resulted in behavioural changes in zebrafish that are relevant to core domains often altered in human psychiatric disorders (such as hyperactivity, impulsivity, reduced habituation and heightened arousal) and suggest that adaptation occurs as the animal develops.

### rbfox1 is expressed in the developing neural and cardiac tissues and in adult brain regions involved in stress, social and emotional behaviours, reward and learning

Consistent with our behavioural results and as seen previously2,3, when we assessed *rbfox1* mRNA distribution in *rbfox1*^*+/+*^ larvae and adults (Tübingen strain), we found that *rbfox1* was expressed in regions of the brain involved in the response to stress, in social and emotional behaviour, and in reward and learning (Figure 1A-B) in agreement with data in rodents36 and humans37.

In wild type larvae *rbfox1* is expressed in the spinal cord and hindbrain lateral neurons (Figure 1A, 28 hpf), whereas at later developmental stages, *rbfox1* expression was detected in the mid- and hindbrain (Figure 1A, 2-5 dpf) and in the heart (Figure 1A, 5dpf) as described previously38.

In adult fish, *rbfox1* is expressed along the entire whole rostro-caudal brain axis. In the forebrain, *rbfox1* was detected in the glomerular (GL), internal (ICL), external (ECL) cellular layers of the olfactory bulbs (Figure 1B a, a’). More caudally, *rbfox1* expression was detected in the medial zone of the dorsal telencephalic area (Dm), and in the dorsal (Vd), lateral (Vl) and ventral (Vv) nuclei of the ventral telencephalic area (Figure 1B b). In the diencephalon, *rbfox1* expression was detected in the ventral habenular nucleus (HaV), in the anterior (A) and ventromedial (VM) thalamic nuclei (Figure 1B c). In the midbrain, *rbfox1* was detected in the posterior part of the parvocellular preoptic nucleus (PPp, Figure 1B d), in the ventral zone of the periventricular hypothalamus (Hv, Figure 1B d, e’), in the periventricular gray zone (PGZ,) and in the central zone (CZ) of the optic tectum (Figure 1B e’), in the torus longitudinalis (TL) (Figure 1B e, e’, g), in the periventricular nucleus of posterior tuberculum (TPp), in the anterior tuberal nucleus (ATN, Figure 2B e, e’’), in the posterior tuberal nucleus of the hypothalamus (PTN, Figure 1B f) and in the paraventricular organ (PVO, Figure 1B f). In the hindbrain, rbfox1 was observed in the lateral division of the valvula cerebelli (Val) (Figure 1B g).

**Figure 2.**
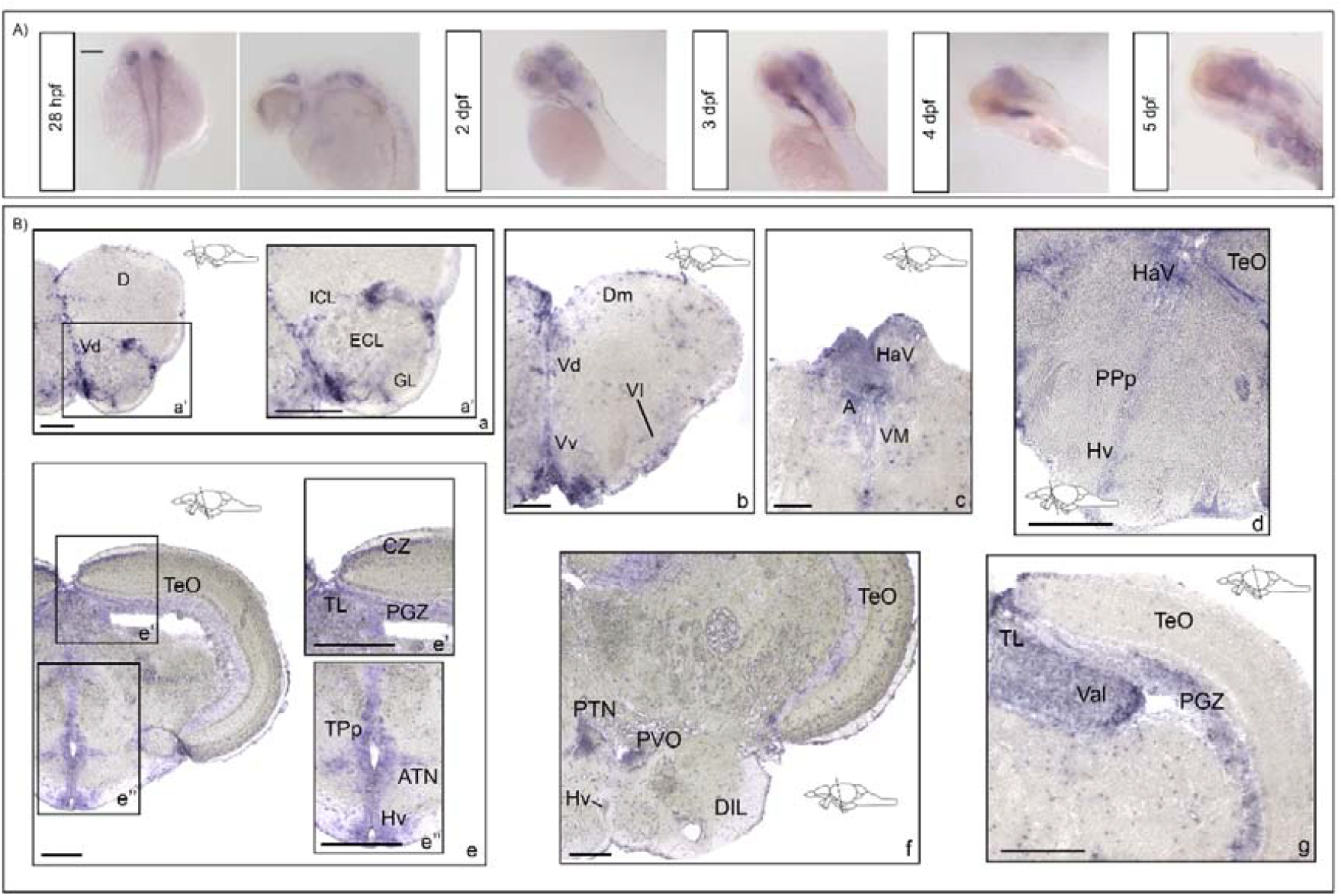
*rbfox1* is expressed in regions of the brain associated with stress response, social and emotional behaviour, reward and learning. **A)** *rbfox1* whole mount *in situ* hybridization (ISH) on zebrafish larvae at 28 hours post fertilization and 3-4-5 days post fertilization. **B)** *rbfox1* ISH on adult zebrafish (a d) forebrain and (e-g) mid-/hind-brain transverse sections. Black boxes represent the region of the brain showed in higher magnification panels. Schematic depictions of the lateral view of the zebrafish brain indicate position of levels illustrated by ISH. Scale bars: 200 µm in A), 100 µm in B) a-c, e, f); 50 µm in B) a’, d, e’, e’’, g. Abbreviations: A, anterior tuberal nucleus; ATN, anterior thalamic nucleus; CZ, central zone of the TeO; D, dorsal area of dorsal telencephalon; DIL, diffuse nucleus of the inferior lobe of the hypothalamus; Dm, dorsal area of medial telencephalon; ECL, external cellular layer; GL, glomerular layer; HaV, ventral area of the habenula; Hv, ventral hypothalamus; ICL, internal cellular layer; PGZ, periventricular gray zone; PPp, parvocellular preoptic nucleus, posterior part; PTN, posterior tuberal nucleus of the hypothalamus; PVO, paraventricular organ; TeO, optic tectum; TL, longitudinal tori; TPp, periventricular nucleus of the posterior tuberculum; Val, valvula cerebelli; Vd, ventral area of dorsal telencephalon; Vl, central area of lateral telencephalon; VM, ventromedial nucleus; Vv, ventral area of ventral telencephalon.

### rbfox1 mutant larvae show elevated cortisol levels and altered expression of *crhb, nr3c2*, bdnf and trkb2

As RBFOX1 has been linked to regulation of HPA axis and *BDNF/TRKB* gene expression4,15,16, we examined the expression of components of the HPI axis and of *bdnf/trkb2*. In parallel, we measured whole-body cortisol levels, in resting physiological conditions, at both larval and adult stages to assess physiological outcomes. Since we did not observe significant differences in behaviour between wild type and heterozygous animals, we employed wild type and homozygous animals only.

We performed qPCR experiments in 5dpf zebrafish larvae to assess changes in the expression levels of the HPI axis markers corticotropin releasing hormone (*crhb*), mineralocorticoid receptor (*nr3c2*), glucocorticoid receptor (nr3c1), corticotropin releasing hormone receptor 1 (*crhr1*), corticotropin releasing hormone receptor 2 (*crhr2*), mineralocorticoid receptor 1 (*mc1r*), proopiomelanocortin a (*pomca*), Krüppel-like factor 9 (*klf9*), as well as of *bdnf* and *trkb2*. In teleosts, the duplication of the genome gave rise to two *CRH* genes, *crha* and *crhb*, and two *TRKB* genes, *trkb1* and *trkb2*39,40. Among these, *crhb* and *trkb2* are most commonly referred to as the functional orthologues of mammalian *CRH* and TRKB respectively, whereas *crha* and *trkb1* function appear more divergent and remain less well understood 39,40. Zebrafish also possess two pomc paralogues (*pomca* and *pomcb*) but only *pomca* is functionally relevant for the activation of the HPI axis41, and six melanocortin receptors (*mc1r, mc2r, mc3r, mc4r, mc5ra, mc5rb*). Among these, we selected *mc1r* as representative of melanocortin signalling. Then, unlike other teleosts, zebrafish possess a single copy of the *MR* (*nr3c2*), GR (*nr3c1*) and *KLF9* (*klf9*) genes, and a single copy of the BDNF gene (*bdnf*)40,42. In mammals, several TRKB splicing isoforms are present, but the most abundant ones are the full-length (TRKB.FL/*TK+*) and the truncated (TRKB.T1/*TK-*), this latter lacking the catalytic tyrosine kinase (TK) domain43. As in zebrafish the presence of both *trkb2* full-length and truncated forms has been demonstrated44, here we used a pair of *trkb2 TK+/TK-* common primers, and another pair targeting only *TK+* to distinguish effects on the expression of the two isoforms.

In *rbfox1*^*19del/19del*^ larvae, inwe observed a significant up-regulation of *bdnf* (p < 0.05) and *TK+* (p < 0.05), and a significant down-regulation of *TK-* (p < 0.05) expression levels (Figure 3A). Regarding HPI axis, in *rbfox1*^*19del/19del*^ larvae we found a significant up-regulation of *crhb* (p < 0.0001) and a significant up-regulation of *nr3c2* (p < 0.01) (Figure 2B). We observed no significant changes in the expression levels of the other genes examined (p > 0.05) (Figure 3B).

**Figure 3.**
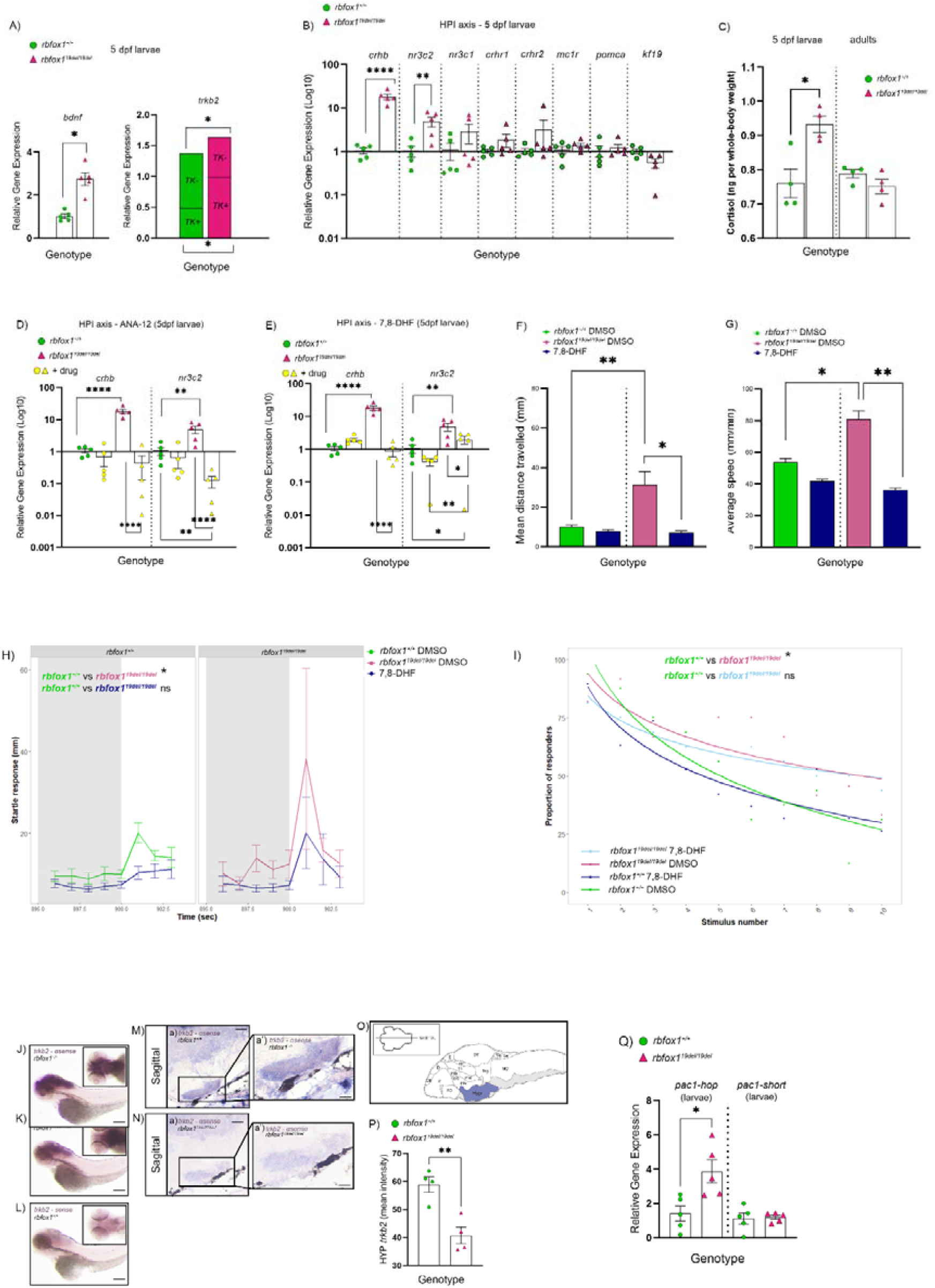
*rbfox1* LoF disrupts zebrafish larvae HPI axis, *bdnf/trkb2* pathway and *pac1a* expression. TRKB modulation restore HPI gene expression and behaviour. **A)** Expression levels of bdnf and trkb2 full-length (*TK+*) and truncated (*TK-*) in 5 days post fertilisation (dpf) larvae (*rbfox1+/+* and *rbfox1*^*19del/19del*^). trkb2 expression is shown in a stacked bar format, where the directly measured *TK+* (long form) is shown as the lower portion and the inferred *TK-* (short form) is shown on top. **B)** Expression levels of HPI axis genes *crhb, nr3c2, nr3c1, crhr1, crhr2, mc1r, pomca* and *kf19* in 5dpf larvae (rbfox1*+/+* and *rbfox1*^*19del/19del*^). **C)** Whole-body cortisol measurement of 5dpf larvae (left panel) and adult fish (right panel) in resting physiological conditions. Cortisol values are normalised per whole-body homogenate (g) (*rbfox1+/+* and *rbfox1*^*19del/19del*^). **D-E)** Expression levels of *crhb* and nr3c2 in presence or absence of the TRKB selective **D)** antagonist ANA-12 and **E)** agonist 7,8-DHF in 5dpf larvae (*rbfox1+/+* and *rbfox1*^*19del/19del*^). **F-I)** Behavioural assays in presence and absence of the TRKB agonist 7,8-DHF in rbfox1 larvae (*rbfox1+/+* and rbfox1^*19del/19del*^): **F)** mean distance travelled, **G)** average speed, **H)** startle response to white light stimulus and **I)** Proportion of responders over time in the response and habituation to acoustic startle assay. **J-N)** Whole mount in situ hybridisation (ISH) for trkb2 anti-sense riboprobe in **J)** *rbfox1+/+* and **K)** *rbfox1*^*19del/19del*^ and **L)** trkb2 sense riboprobe in *rbfox1+/+* 5dpf larvae (lateral view in the main boxes and dorsal view in the smaller boxes on the top right corner). **M, N)** Sagittal cryosection of trkb2 ISH in **M)** *rbfox1+/+* and **N)** *rbfox1*^*19del/19del*^ 5 dpf larvae. Black boxes in **M-a)** and **N-a)** represent the region of the brain showed in higher magnification panels in **M-a’)** and **N-a’)**. Scale bars: 200 µm in J), K) and L); 100 µm in M-a) and N-a); 50 µm in **M-a’)** and **N-a’). O)** Schematic depiction (sagittal) of zebrafish larval brain indicating position of levels illustrated by ISH on sagittal cryosections. **P)** trkb2 ISH intensity mean in the hypothalamus of 5dpf zebrafish larvae, *rbfox1+/+* versus *rbfox1*^*19del/19del*^. N = 4 larvae x genotype. **Q)** Expression levels of pac1-hop and pac1-short in *rbfox1+/+* and *rbfox1*^*19del/19del*^ 5 dpf larvae. For qPCR experiments, reference genes were actin – β 2 (*actb2*) and ribosomal protein L13a (*rpl13a*). Each green dot/pink triangle in A-D, K-L) represents a pool of 15 larval heads (eyes and jaw removed), while yellow ones represent larvae exposed to TRKB drugs. Where indicated, we used Log_10_ transformation to normalize the data facilitating a clearer visualisation of trends within the dataset. All larvae employed were progeny of *rbfox1+/19del* in-cross and were genotyped after experiments and prior to data analysis. For cortisol measurement (G), each dot/triangle represent a pool of 12 larvae in the left panel (larvae) and a single whole zebrafish in the right panel (adult fish). In all graphs: bars represent standard error of the mean (SEM); * p < 0.05; ** p < 0.01; **** p < 0.0001.

To assess rbfox1 LoF effects on zebrafish larvae during early developmental stages, we performed *crhb, bdnf* and *trkb2* qPCR experiments also in 3dpf larvae. Similarly to 5dpf larvae, in 3dpf *rbfox1*^*19del/19del*^ larvae we observed a significant up-regulation of *crhb* (p < 0.01), bdnf (p < 0.05) and *TK+* (p < 0.05), and a significant down-regulation of *TK-* (p < 0.05) (Supplementary Figure 4A).

We measured whole-body cortisol levels under resting physiological conditions in both larvae and adult zebrafish to evaluate the physiological impact of *rbfox1* loss and observed changes in *crhb* expression. We found a significant increase in cortisol levels in *rbfox1*^*19del/19del*^ larvae (p < 0.05) (Figure 3C). In contrast, we observed no significant differences in adult fish (p > 0.05) (Figure 3C).These findings showed that *rbfox1* LoF led to alterations in stress-related gene expression and cortisol levels in larvae, in line with our behavioural data and implicating *RBFOX1* as a critical regulator of stress response mechanisms.

### Pharmacological treatment targeting TrkB signalling restores *crhb* and *nr3c2* gene expression and disrupted hyperactivity, startle response and habituation in rbfox1 LoF larvae

To assess whether molecular and behavioural changes seen in *rbfox1* LoF larvae were mediated by bdnf/trkb2 signalling, we repeated qPCRs and behavioural experiments in 5dpf zebrafish larvae following chronic exposure (from 5 h to 5dpf) to the TRKB agonist 7,8-DHF or antagonist ANA-12. As in our line *rbfox1* expression was knocked down throughout development, we initiated TRKB drug treatment from gastrulation.

In *rbfox1*^*+/+*^ larvae, neither TRKB agonist 7,8-DHF or antagonist ANA-12 had a significant effect on *crhb* and *nr3c2* gene expression (p > 0.05) (Figure 3D, E). In *rbfox1*^*19del/19del*^ larvae, chronic exposure to either TRKB agonist or antagonist significantly reduced *crhb* and *nr3c2* expression levels relative to untreated mutant larvae (p < 0.001), restoring gene expression to levels comparable with wild type controls (p > 0.05) (Figure 3D, E). Following identification and removal of outliers (1 outlier in the *rbfox1*^*+/+*^ + ANA-12 group, 1 outlier in the *rbfox1*^*+/+*^ + 7,8-DHF group and 1 outlier in the *rbfox1*^*19del/19del*^+ 7,8-DHF group), we observed a significant down-regulation of *nr3c2* expression levels in *rbfox1*^*19del/19del*^ + ANA-12 versus *rbfox1*^*+/+*^ un-exposed controls (p < 0.01) (Figure 2D) and a significant up-regulation of *nr3c2* expression levels in *rbfox1*^*19del/19del*^+ 7,8-DHF versus both exposed (p < 0.01) and un-exposed (p < 0.05) controls (Figure 3D, EE). Interestingly, while having the same effect on *crhb* and *nr3c3* expression, ANA-12 and 7,8-DHF had different outcomes on *bdnf* expression. Chronic treatment with ANA-12 reduced *bdnf* expression in *rbfox1*^*+/+*^ (p < 0.05) but had no effect on *rbfox1*^*19del/19del*^. On the other hand, chronic treatment with 7,8-DHF increased bdnf levels in both *rbfox1*^*+/+*^ and *rbfox1*^*19del/19del*^ larvae (Supplementary Figure C).

Then, we examined the effects of ANA-12 and 7,8-DHF on behaviour to assess which behavioural alterations observed in *rbfox1* mutant were influenced by bdnf/trkb2 signalling. In presence of 7,8-DHF we found a significant two-way interaction between genotype and drug on distance travelled [Effect of genotype by drug: χ2(2) = 47.3782, p < 0.05] and on swimming velocity [Effect of genotype by drug: χ2(2) = 31.3951, p < 0.05] such that treatment with 7,8-DHF restored hyperactive behaviour observed in *rbfox1*^*19del/19del*^ larvae (p < 0.01) (Figure 3F, G). Further, following chronic exposure to 7,8-DHF there was no significant difference between *rbfox1*^*19del/19del*^ and *rbfox1*^*+/+*^ larvae in response to startle with white light (Figure 3H) or in the proportion of responders to the acoustic startle (p > 0.05) (Figure 3 I). ANA-12 had no significant effects on *rbfox1*^*19del/19del*^ behavioural alterations.

As the hypothalamus is the primary player in the stress response and Rbfox1 had only previously been shown to affect TRKB expression within the hippocampus16, we used ISH to determine whether trkb2 mRNA expression was reduced within the hypothalamic area. In agreement with previous findings45, in *rbfox1*^*+/+*^ we found that *trkb2* is widely expressed in the brain of 5dpf larvae, whereas in *rbfox1*^*19del/19del*^ larvae, consistently with our qPCR experiments, we found a significant overall reduction of trkb2 mRNA in the whole brain (Figure 3J-L), including in the hypothalamus (p < 0.01) (Figure 3M, N, P, Supplementary Figure 5).Our findings showed that the dysregulation in *crhb* and *nr3c2* expression caused by *rbfox1* LoF was prevented by TRKB agonists and antagonists. Further, we showed that hyperactivity, startle hypersensitivity and heightened proportion of responders to the acoustic startles observed in *rbfox1* mutants are at least partially mediated by disrupted bdnf/trkb2 signalling.

### rbfox1 LoF alters pac1a expression levels of zebrafish larvae

As RBFOX proteins have been shown to be able to regulate *Pac1* alternative splicing to include the *hop* cassette7 and PAC1 has been shown to regulate *Bdnf* transcription and potentiate TRKB activity17, we examined expression levels of *pac1a*, the zebrafish homologue of the mammalian *PAC1*, in *rbfox1*^*+/+*^ and *rbfox1*^*19del/19del*^ 5dpf larvae by qPCR.

In mammals there are several PAC1 isoforms and their role in the regulation of stress is poorly understood9. In the brain, the predominant isoforms are *PAC1-hop* (long isoform) and *PAC1-short*9. Zebrafish possess two *pac1* genes, *pac1a* and *pac1b*, but only *pac1a* contains the hop cassette9. PAC1-short enhances *CRH* transcription, while PAC1-hop reduces *CRH* synthesis during late stress recovery phase4. We measured expression levels of both *pac1a-short* and -*hop* isoforms and we observed no differences in *pac1a-short* expression, but we found a significant up-regulation of *pac1a-hop* in *rbfox1*^*19del/19del*^ mutant larvae (p < 0.05) (Figure 3Q).

This finding shows that elevated pac1a-hop/Pac1a-hop alone is not sufficient to counteract *crhb* increase and strengthens previous data7 suggesting that RBFOX1 is not the main regulator (or at least not the sole regulator of) *PAC1* alternative splicing..

### rbfox1 mutants undergo adaptive mechanisms and allostatic overload during development

The HPA axis possesses a vital role in the maintenance of allostasis, the process by which the body achieves stability in response to stress or environmental challenges. Dysregulation of the HPA axis often leads to disrupted allostasis during later life, also termed as allostatic load (i.e., the physiological consequence resulting from the cumulative “wear and tear” of the body in response to chronic stress)22. As we observed dysregulation of the HPI axis gene expression in *rbfox1* mutant larvae and altered behavioural responses in *rbfox1* mutant larvae but not in adults, we assessed expression of HPI axis components in adult zebrafish, in the presence and absence of acute stress (NTD), to explore the possibility of adaptation that may contribute to differences in allostatic load.

We first measured *crhb, nr3c2, nr3c1, bdnf* and *trkb2* mRNA expression levels. In physiological resting conditions, we observed no differences in HPI nor *bdnf/trkb2* expression levels (p > 0.05) (Figure 4A, B). However, after stress exposure, regarding the HPI axis, in *rbfox1*^*19del/19del*^ adults we observed the same dysregulation seen in rbfox1 LoF larvae: we found significant up-regulation of *crhb* (p < 0.001) and *mr* (p < 0.0001) and no changes in *nr3c1* expression levels (p > 0.05) (Figure 4A), and significant up-regulation of *bdnf* (p < 0.01) and *TK+* (p < 0.05), and significant down-regulation of *TK-* (p < 0.05) (Figure 4B, C).

**Figure 4.**
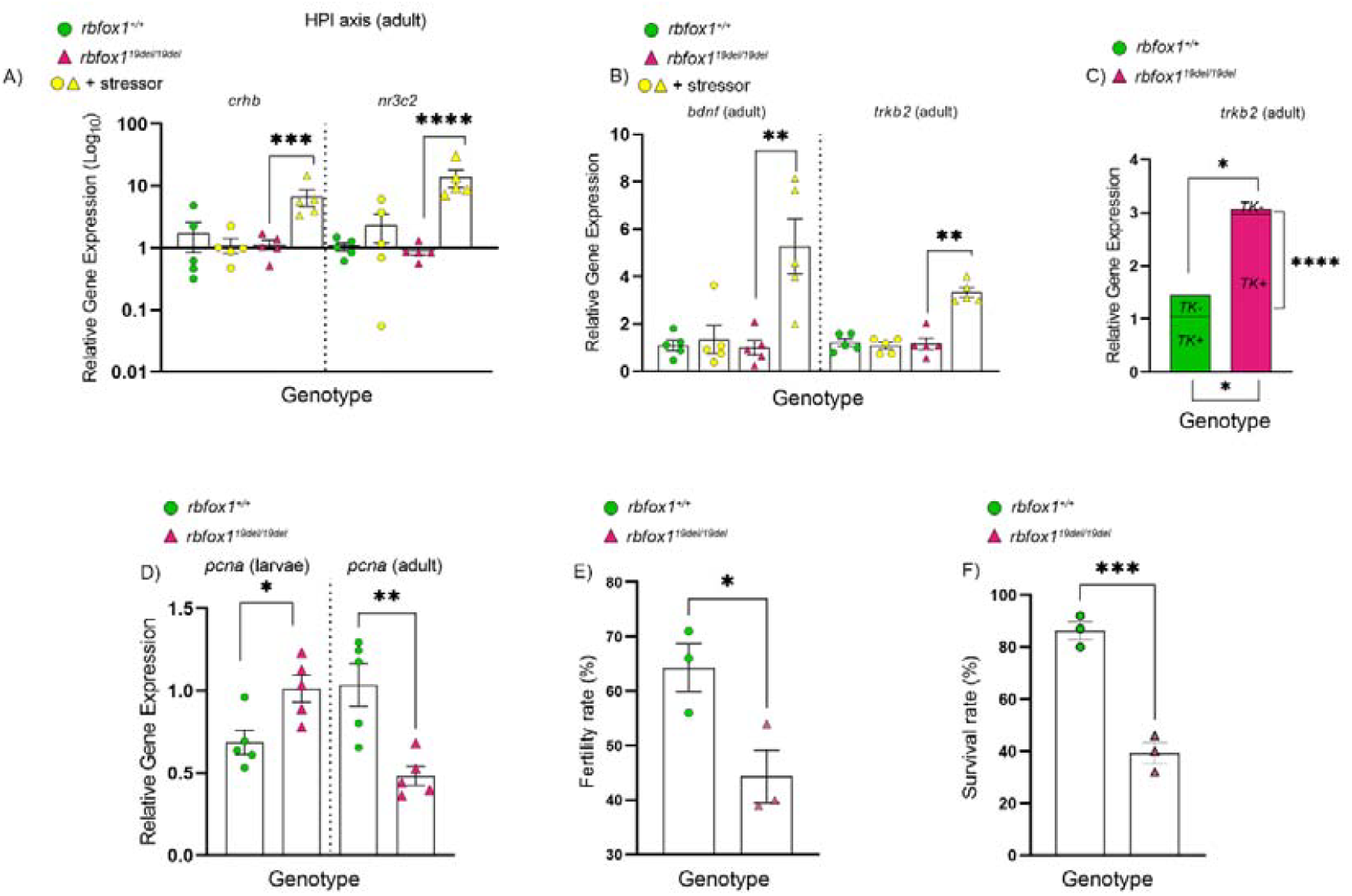
*rbfox1* mutants undergo adaptive mechanisms and allostatic overload during development. Expression levels of **A)** HPI axis genes corticotropin releasing hormone b (*crhb*), mineral corticoid receptor (*mr*) and glucocorticoid receptor (*gr*) and of **B)** *bdnf* and *trkb2* truncated/full-length (*TK-/TK+*) (common primers) in adult zebrafish brain (*rbfox1+/+* and *rbfox1*^*19del/19del*^) in normal resting conditions and after exposure to a stressor (novel tank diving). **C)** Expression levels of the *trkb2 TK+* and *TK-* in adult zebrafish brain (*rbfox1+/+* and *rbfox1*^*19del/19del*^) after exposure to a stressor.). trkb2 expression is shown in a stacked bar format, where the directly measured *TK+* (long form) is shown as the lower portion and the inferred *TK-* (short form) is shown on top. **D)** Expression levels of proliferating cell nuclear antigen (*pcna*) in 5 days post fertilisation zebrafish larvae and adult zebrafish brain (*rbfox1+/+* and r*bfox1*^*19del/19del*^) in normal resting conditions. **E)** Fertility rate and **F)** survival rate of *rbfox1+/+* and *rbfox1*^19del/19del^. Each green dot/pink triangle in A-B) represents a single adult *rbfox1+/+* or *rbfox1*^*19del/19del*^ brain under resting physiological conditions respectively, while yellow dots/triangles represent *rbfox1+/+* or *rbfox1*^*19del/19del*^ brain after stress exposure respectively. In B) each green dot/pink triangle represents single adult *rbfox1+/+* or *rbfox1*^*19del/19del*^ brain after stress exposure respectively. In D) for larvae each green dot/pink triangle represents a pool of *rbfox1*^*+/+*^ or *rbfox1*^*19del/19del*^ 15 larval heads (eyes and jaw removed) respectively; for adults each green dot/pink triangle represents a single adult *rbfox1+/+* or *rbfox1*^*19del/19del*^ brain respectively. In E) each dot/triangle represents average fertility of 3-5 trios (1 male and 2 females) assessed over 2-3 petri dish (50 embryos per dish) per trio. Each trio belonged to a different tank (for each genotype for each batch) to avoid tank effect). In F) each dot/triangle represents the percentage of survival of a single fish stock comprising 50 larvae. For qPCR experiments, reference genes were actin – β 2 (*actb2*) and ribosomal protein L13a (*rpl13a*). Where indicated, we used Log_10_ transformation to normalize the data facilitating a clearer visualization of trends within the dataset. All larvae employed were progeny of *rbfox1*^*+/19del*^ in-cross and were genotyped after experiments and prior to data analysis. In all graphs: bars represent standard error of the mean (SEM); * p < 0.05; ** p < 0.01; *** p < 0.001; **** p < 0.0001.

Excess glucocorticoid (GC) exposure during early developmental stages or early life stress (ELS) can lead to vulnerability to allostatic overload in the long-term19,22. One of the effects caused by GC-or ELS-induced allostatic overload is reduced cell proliferation during adulthood19. This effect has been suggested to be developmentally dynamic, since it is often preceded by increased cell proliferation during early stages19. Further, BDNF signalling has been shown to regulate neural stem cell proliferation through *TK-*, suggesting that *rbfox1* LoF fish showing altered levels of *TK-* expression, may show altered proliferation across the life course46. We therefore assessed the rate of proliferation at both larval and adult stages using qPCR for proliferating cell nuclear antigen (pcna). We observed a significant up-regulation of pcna expression levels in 5dpf *rbfox1*^*19del/19del*^ zebrafish larvae (p < 0.05) and a significant down-regulation of *pcna* expression levels in the brains of adult *rbfox1*^*19del/19del*^ fish (p < 0.01) when compared to *rbfox1*^*+/+*^ siblings (Figure 4D).

In adult zebrafish, other effects caused by GC-induced allostatic overload include reduced fertility and survival rates19. Therefore, we assessed fertility and survival of *rbfox1*^*19del/19del*^ zebrafish and *rbfox1*^*+/+*^ siblings. We found that both fertility (p < 0.05) and survival (p < 0.001) rates of *rbfox1*^*19del/19del*^ fish were significantly reduced compared to rbfox1*+/+* siblings (Figure 4E, F).

Consistent with our findings (disrupted HPI axis gene expression and increased cortisol in *rbfox1* mutant larvae under resting physiological conditions and in *rbfox1* adult mutant only following stress exposure), these results suggest that *rbfox1* mutants may engage compensatory mechanisms during development similar to those triggered by early-life GC exposure. While these mechanisms may transiently restore homeostasis, they likely lead to enduring alterations in stress responsivity and neurodevelopmental trajectories.

## DISCUSSION

In this study we generated a CRISPR-Cas9 LoF *rbfox1* zebrafish line (*rbfox1*^*19del*^) to investigate the mechanisms by which *RBFOX1* LoF increases susceptibility to psychiatric disorders.

Allostatic load has been linked with several cognitive disorders including depression, schizophrenia, anxiety and PTSD47. HPA axis overactivity is a key factor in the onset of the allostatic load22. RBFOX1 has been shown to influence expression of stress-related genes such as PAC1 and BDNF/TRKB. Therefore, alterations in RBFOX1 function may increase psychiatric disorder vulnerability through alterations in stress response systems, such as the HPA axis, pre-disposing to allostatic overload vulnerability in later life (Figure 5).

**Figure 5.**
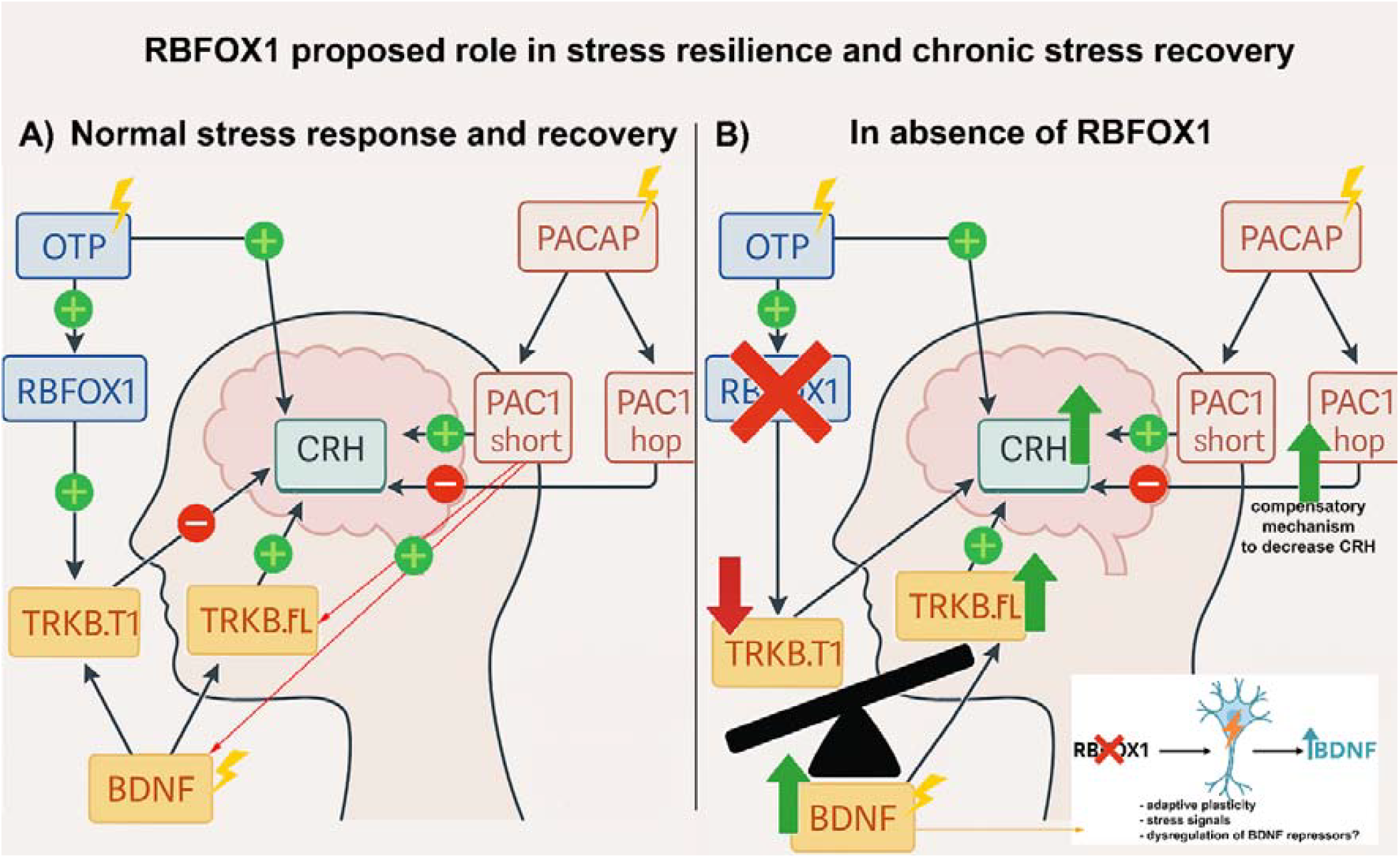
RBFOX1 role in stress resilience and chronic stress recovery: proposed mechanism of action. **A)** In response to stressful challenges (yellow flash), BDNF expression levels increase, stimulating *CRH* transcription via TRKB.FL/TK+ (yellow pathway). At the same time, stress triggers also *PACAP* transcription (red pathway) and OTP-mediated transcription of *RBFOX1* and *CRH* (blue pathway). *PACAP* stimulates *CRH* transcription via PAC1-short and inhibits *CRH* transcription via PAC1-hop. PACAP binding to PAC1-short also increases *BDNF* transcription (red line) and enhances TRKB.FL/TK+ activity (red line). RBFOX1 functions as a “switch” maintaining the balance between full-length/truncated TRKB isoforms (promoting TRKB mRNA stability and/or alternative splicing): during the stress recovery phase, *RBFOX1* switches the balances in support of *TRKB*.*T1/TK*-isoform, decreasing CRH levels and turning off the HPA axis. **B)** *RBFOX1* deletion leads to an increase in *BDNF* expression levels (yellow arrow and white box) and an unbalance between *TRKB*.*FL/TK+* (increased) and *TRKB*.*T1/TK-* (decreased) isoforms. Such dysregulation ultimately leads to an increase in CRH transcription (and possible hyperactivation of the HPA axis), and increase in PAC1-hop levels as possible compensatory mechanism. Our data suggest that RBFOX1-mediated regulation of *TRKB*.*T1/TK-* mRNA stability and/or alternative splicing is necessary to promote neuroplasticity and stress resilience.

Similarly to Rbfox1 deficient mice6 and as seen in previous studies employing *rbfox1* mutant zebrafish2,23, we found that *rbfox1*^*19del/19del*^ mutants were hyperactive and impulsive. These findings are consistent with studies implicating common and rare *RBFOX1* genetic variants as risk factors for psychiatric disorders like ASD, ADHD, schizophrenia and anxiety/stress disorders5,6,48. At larval stages *rbfox1*^*19del/19del*^ were hyperactive and had increased burst swimming, indicative of increased impulsivity. In the response and habituation to acoustic startle assay, *rbfox1*^*19del/19del*^ larvae startled more and had a higher response rate over time compared to *rbfox1*^*+/+*^ siblings, consistent with previous findings in rbfox1 mutant zebrafish30. In a forced light-dark assay, although baseline hyperactivity of *rbfox1*^*19del/19del*^ larvae makes interpretation difficult, *rbfox1*^*19del/19del*^ larvae showed an increased startle response upon dark to light transition, consistent with heightened anxiety. These results suggest that *rbfox1* LoF leads to heightened arousal, aligning with studies linking *RBFOX1* to mood and anxiety disorders, including PTSD49. Interestingly, *rbfox1*^*19del/19del*^ adults showed reduced bottom dwelling only during the first minute of the novel tank diving assay (an adult measure of anxiety-like behaviour) while there were no significant differences during the remaining 4 minutes of the assay This latter result suggests behavioural adaptation possibly coupled with increased impulsivity or impaired threat assessment (decreased bottom dwelling during the first minute of assay. This aligns with the broader *rbfox1*^*19del/19del*^ behavioural profile (hyperactivity, heightened startle and poor habituation at larval stages) indicative of dysregulated arousal rather than reduced anxiety. Similarly, in the 5-CSRTT we found that *rbfox1*^*19del/19del*^ were more impulsive than rbfox1*+/+* siblings, consistent with RBFOX1 variants being associated with impulsivity and related disorders such as ADHD and ASD6.

In *rbfox1* LoF larvae, under resting physiological conditions, we observed increased cortisol levels and dysregulation of *crhb* and *nr3c2*, both key components of fish HPI axis, which is analogous to the HPA axis in mammals. The HPA axis is crucial for managing stress, with *CRH* triggering cortisol secretion, affecting metabolism, immunity, and behaviour50,51. In zebrafish larvae under resting physiological conditions, we found increased cortisol, *crhb* and *nr3c2*, with no changes in the expression levels of the other HPI axis genes assessed (nr3c1, *crh*r1, *crh*r2, mc1r, pomca, kf19). In adults, although resting levels were unchanged, rbfox1 LoF showed an exaggerated molecular response to stress. Notably, both *CRH* and MR have important roles in the brain beyond their canonical endocrine functions via the HPA axis. *CRH* regulates anxiety and arousal acting as a neuromodulator in limbic regions such as the amygdala and hippocampus52. Similarly, hippocampal MRs play a key role in early stress responses by supporting cognitive appraisal of novel situations and promoting behavioural flexibility, thus shaping the initial reaction to stress53. Elevated cortisol and *crhb* levels as seen in rbfox1^19del/19del^ larvae under resting physiological conditions, suggest a chronic anxiety state. Increased cortisol, *CRH* and MR levels are linked to mood disorders like depression and PTSD54–57. These findings support a role for RBFOX1 in *CRH* regulation and suggest that RBFOX1 LoF may contribute to mood disorders and stress resilience via HPA axis dysregulation and/or non-canonical brain-specific effects of *CRH* and MR.

BDNF and TRKB are also key regulators of the stress response58. BDNF levels often increase during stress to promote neuronal survival and plasticity, buffering negative effects of stress on the brain via TRKB10,11,13,58–60. Dysfunction of BDNF/TRKB signalling is linked to several stress-related disorders13. The increased bdnf expression we observed in rbfox1^19del/19del^ larvae may suggest a compensatory response to cope with *crhb* increase and maintain homeostasis. Given BDNF’s established role in modulating neuronal plasticity and stress adaptation, its upregulation may serve to stabilise or buffer hypothalamic circuits disrupted by excessive *crhb* transcription58. This response could represent an intrinsic attempt to mitigate possible overactivation of the HPI axis or the reduced activity within the trkb2 signalling (a suggestion supported by finding that incubation in the TRKB agonist 7,8-DHF rescues the behaviour as well as the expression of *crhb*) and maintain functional equilibrium in the face of impaired regulatory input caused by *rbfox1* loss. Alternatively, the observed increase in *bdnf* expression could result from either dysregulation of BDNF repressors or an imbalance in TRKB isoform expression caused by RBFOX1 loss, where upregulation of the full-length isoform may enhance BDNF expression through a positive feedback mechanism40,61. At 3 and 5dpf, in *rbfox1*^*+/+*^ larvae we detected both *trkb2* truncated (*TK-*) and full-length (*TK+*) forms, and in *rbfox1*^*19del/19del*^ larvae we observed a significant down-regulation of *TK-* and a significant up-regulation of *TK+*, which is predicted to lead to an increased activation of TRKB signalling, consistent with the increased *crhb* expression as seen here62. Studies in rodents showed that *Rbfox1* up-regulation led to increased TrkB.T1 (*TK-*) expression in the hippocampus and no changes were observed following *Rbfox1* deletion16. In contrast, we observed down-regulation of *trkb2 TK-* expression in *rbfox1*^*19del/19del*^ larvae in resting physiological conditions. The difference between these studies may be due to i) differences in the tissue examined (zebrafish larvae whole head versus rodent hippocampus), ii) the developmental stage of the animals (larval versus adult stage), iii) the genotype of the animals studied (*rbfox1*^*19del/19del*^ in our study versus *Rbfox1*^*+/-*^ in the study in rodents), or iv) presence of compensatory mechanisms (in the hippocampal mouse model *Rbfox1* knockdown resulted in up-regulation of *Rbfox2*). As for this latter hypothesis, as in previous studies in zebrafish2, we did not see any changes in *rbfox2* expression upon *rbfox1* LoF (Supplementary Figure4) suggesting that this compensatory mechanism does not occur in larval fish.

Consistent with the possibility that *RBFOX1* LoF may contribute to dysregulation of the HPA axis through a direct or indirect effect on BDNF/TRKB pathway, chronic exposure to TRKB agonist (7,8-DHF) or antagonist (ANA-12) restored HPI axis gene expression to that seen in *rbfox1*^*+/+*^ larvae. ANA-12 is a selective non-competitive antagonist of TRKB exerting central TRKB blockade and producing rapid and long-lasting anxiolytic and antidepressant effects63. Mice treated with ANA-12 showed reduced anxiety-like behaviour63 and in stressed rats, ANA-12 blocked *Crh* increase in the hypothalamus and amygdala64. 7,8-DHF is a potent selective agonist of TRKB used in the treatment of several disorders, including depression and schizophrenia, and has been shown to enhance memory consolidation and emotional learning in healthy rodents65. In a similar fashion to the effects of ANA-12 seen in rats (blocking *Crh* increase)57, after chronic treatment with ANA-12, we observed that the expression of HPI axis genes in *rbfox1*^*19del/19del*^ larvae was restored to *rbfox1*^*+/+*^ levels. 66 Interestingly, ANA-12, despite restoring *crhb* and *nr3c2* levels to wild type, did not rescue *rbfox1*^*19del/19del*^ larvae behaviour, indicating that blockade of TRKB can normalise some aspects of HPI axis output but is insufficient to correct the behavioural phenotype (perhaps because it further disrupts necessary TRKB-mediated signalling in the brain). In contrast, 7,8-DHF, normalises *crhb* and *nr3c2* expression and rescues some behavioural aspects altered in *rbfox1*^*19del/19del*^ larvae (i.e., hyperactivity, startle response to white light stimulus and proportion of fish responding to acoustic stimuli), suggesting that enhancing TRKB signalling through the full-length receptor may restore functional signalling balance and downstream pathways (e.g., synaptic plasticity, habituation mechanisms). Finally, the fact that 7,8-DHF increases *bdnf* expression in *rbfox1*^*19del/19del*^ larvae while ANA-12 has no effect suggests a potential positive feedback loop whereby TRKB activation enhances BDNF expression, helping restore adaptive neuroendocrine and behavioral responses. Future studies employing acute or temporally restricted treatments (e.g., 3–5 dpf), or employing different TRKB agonist/antagonist, will help delineate the specificity of TRKB signalling effects. Interestingly, *rbfox1*^*19del/19del*^ larvae were differentially sensitive to the effect of TRKB drugs on *nr3c2* expression, most likely due to the unbalanced *TK+/TK-* levels. These findings warrant further investigations. It is of note that neither TRKB agonist nor antagonist had a significant effect on *crhb* nor *n3cr2* expression in *rbfox1*^*+/+*^ larvae. One possible explanation is that, as the larvae were not exposed to any stressors, any effect on Trkb2 signalling had limited effect on HPI axis activity.

In *rbfox1*^*19del/19del*^ larvae, we also observed up-regulation of *pac1a-hop* with no changes in *pac1a*-*short* levels. Previous studies in zebrafish have shown that Pac1-short gain of function caused persistent *crh* increase, while overexpression of Pac1-hop prevented stress-induced *crh* transcription activation4. These previous studies also suggest that RBFOX1 regulates the alternative splicing of *PAC1* promoting the formation of *PAC1-hop*4. However, the increase in *pac1-hop* seen in our study argues against this suggestion and suggests an adaptive response to the increased expression of *crh* either through *rbfox2*7 (whose expression was not altered by *rbfox1* LoF (Supplementary Figure4)) or through an unknown mechanism. Elevated *pac1a-hop*/Pac1a-hop alone might be not sufficient to counteract *crhb* upregulation, given the concurrent increase in *bdnf* levels and the shift in trkb2 isoform expression (upregulation of the excitatory long form and downregulation of the inhibitory short form). This interpretation is further supported by the decreased in *crhb* levels observed following chronic TRKB modulation. Another option is that the upregulation of *pac1a-hop* mRNA may not reflect increased functional Pac1a-hop receptor activity, particularly in the context of disrupted splicing regulation following rbfox1 loss, suggesting that RBFOX1 might be involved in PAC1 mRNA stability rather than alternative splicing.

In contrast to larvae, differences in HPI axis and *bdnf/trkb2* gene expression and cortisol levels were not seen in adult animals under resting conditions. However, *rbfox1*^*19del/19del*^ adults showed increased *crhb* and *nr3c2* on challenge with a stressor, aligning with the previous suggestion that Rbfox1 is required for the termination of the acute endocrine stress response4 and that chronic stress during development leads to adult HPI axis adaptation, often at a cost in later life (e.g., altered stress responsivity, reduced fertility and survival, decreased *pcna* expression)19,67. This interpretation is consistent with previous work by Eachus et al., which demonstrated that early-life exposure to excess GC can lead to long-term changes in stress-regulatory circuits and adult behavioural dysfunction19. Supporting this, *rbfox1* LoF adults show heightened *crhb, nr3c2* and *bdnf* expression, and imbalance in *trkb2* isoform expression, specifically in response to a stressor (the novel tank diving assay). Together, these findings suggest that while baseline homeostasis may be restored through developmental adaptation, stress responsivity remains dysregulated, pointing to an allostatic shift toward a passive coping style for *rbfox1* mutants68,69. Passive (cautious, risk-averse, high *CRH*) and active (bold, exploratory, low *CRH*) coping styles refer to behavioural strategies that animals employ to respond to stress and that are underpinned by distinct neuroendocrine pathways, providing a valuable framework for interpreting stress susceptibility, resilience and the behavioral consequences of genetic mutations68,69. The shift toward a passive coping style of *rbfox1* mutants support a role for RBFOX1 in shaping stress responsivity and behavioural strategies. In *rbfox1*^*19del/19del*^ adults we observed also reduced fertility and reduced survival rates, two long-term effects often associated with excess GC-exposure/early life stress19. Another long-term effect of excess GC-exposure/early life stress is reduced neural stem cell proliferation in adult animals, often accompanied by increased proliferation at early stages19. Our findings that *rbfox1*^*19del/19del*^ showed increased *pcna* levels at larval staged but reduced *pcna* levels during adulthood, corroborate this latter hypothesis of a dynamic developmental effect of early life stress. Consistent with the *crhb* increase observed in stressed *rbfox1*^*19del/19del*^ adults, and as seen in larvae, after acute stress, *rbfox1*^*19del/19del*^ adults also showed unbalanced trkb2 expression, with up-regulation of *TK+* and down-regulation of *TK-*.

In conclusion, given the conservation between fish HPI axis and the mammalian HPA axis, our data unveils a pivotal new finding in *CRH* regulation, revealing an interplay between RBFOX1 and BDNF/TRKB in the context of chronic stress and stress resilience and suggests that RBFOX1 plays a crucial role in adaptive stress mechanisms. In response to stressful challenges, RBFOX1 functions as a “switch” regulating the balance between short/long isoforms of TRKB receptors: during the stress recovery phase, RBFOX1 switches the balance towards *TRKB*.*T1/TK-* isoform, decreasing *CRH* levels and turning off the HPA axis (Figure 5). In *rbfox1*^*19del/19del*^, this regulatory switch appears disrupted (increased cortisol, *crhb, nr3c2, bdnf*, and full-length *trkb* expression alongside reduced truncated trkb) suggesting a failure to terminate the stress response. Behaviourally, mutants exhibited hyperactivity, impulsivity, heightened arousal, and reduced habituation, phenotypes that parallel core symptoms seen in several human psychiatric disorders. Pharmacological experiments support a functional consequence of this dysregulation. Although both ANA-12 (TRKB antagonist) and 7,8-DHF (TRKB agonist) normalised *crhb* and *nr3c2* levels, only 7,8-DHF rescued behavioural abnormalities. These findings suggest that restoring balanced TRKB signalling (by selectively enhancing the activity of residual inhibitory TRKB.T1 receptors without compromising the beneficial effects mediated by excitatory TRKB.FL receptors) is critical for functional recovery. This supports the idea that an imbalance in TRKB isoform expression, rather than overall receptor activity, may underlie the observed behavioural pathology. The differential effects of these compounds also point to a possible role for positive feedback in BDNF expression via TRKB-FL signalling, which may be essential for behavioral homeostasis. Together these results suggest that *RBFOX1* contributes to the liability of psychiatric disorders through *CRH*, MR and BDNF/TRKB dysregulation, which leads to disrupted development and vulnerability to allostatic overload in later life. The suite of behavioural effects seen in *rbfox1* LoF animals coupled with exaggerated response on exposure to a stressor are reminiscent of reported traits in passive versus active coping styles68 suggesting that RBFOX1 activity may also play a key role in individual differences in coping strategies. Although we show a key role for RBFOX1 in regulation of the HPA axis, consistent with this being a primary mechanism by which *RBFOX1* variants pre-disposes to psychiatric disorders, RBFOX1 regulates the splicing of a wide range of additional genes15 which may also play a part of disease vulnerability.

## METHODS

### Animal maintenance

All fish were maintained in a recirculating system (Tecniplast, UK) with a 14h:10h light/dark cycle and a constant temperature of 28°C. Fish were fed with ZM-400 fry food (Zebrafish Management Ltd.) in the morning and brine shrimps in the afternoon. Breeding was set up in the evening, in sloping breeding tanks (Tecniplast) provided with dividers for timed mating. The following morning, dividers were removed to allow spawning. Eggs were collected in Petri dishes (max 50 eggs/dish). Infertile eggs were removed, and fertile ones were incubated at 28°C. Petri dishes were checked daily to ensure consistent developmental stage across groups. If reared, larvae were moved to the recirculating system at 5 days post fertilization (dpf) and fed as stated above.

All procedures were carried out under license in accordance with the Animals (Scientific Procedures) Act, 1986 and under guidance from the Local Animal Welfare and Ethical Review Board at Queen Mary University of London.

Although our previous studies examined two distinct rbfox1 mutant lines (*rbfox1*^*del19*^ _vs rbfox1_^*sa15940*^)2, we chose to focus on the *rbfox1*^*del19*^ line for the current study due to its more pronounced phenotype.

### Generation of rbfox1 loss of function zebrafish line

The zebrafish rbfox1 loss of function (LoF) mutant line was generated as described previously using Tübingen strain as background 71. CRISPR RNA (crRNA) (Merck) was designed to target *rbfox1* exon 2 (CCCAGTTCGCTCCCCCTCAGAAC, PAM sequence in bold, MwoI recognition site underlined). A 3 µL injection mix containing 1 µL (FC 83 ng/µL) crRNA, 1 µL (FC 83 ng/µL) tracrRNA (Merck, #TRACRRNA05N), 1 µL (FC 1.67 µM) and 1 µL Cas9 protein (New England Biolabs, #M0646) was freshly prepared on the morning of the injection procedure. Then, 1 nL of the injection mix was injected into one-cell stage zebrafish embryos (∼ 100-150 embryos). Injection efficacy was assessed at 24 hours post fertilization (hpf) by polymerase chain reaction (PCR) from genomic DNA (*rbfox1*_Forward, 5⍰-TAATCAAGACGCCCCAGCAC–3⍰; rbfox1_Reverse, 5⍰-GTACTCAGCAGGAATGCCGT-3⍰) followed by MwoI (New England Biolabs, #R0573S) restriction enzyme digestion. Successful injections will introduce indel mutations disrupting the recognition site of the restriction enzyme, preventing this latter from cutting the PCR amplicon. Once reached sexual maturity (at ∼ 3 months of age) injected fish (F0) were outcrossed with wild type to generate F1 embryos. The progeny may carry different mutations due to the mosaic nature of the F0 parents. F1 fish were therefore screened for mutations leading to premature termination codon (PTC) via cloning into pGEM-T Easy vector (Promega, #A1360), followed by transformation, colony PCR, DNA purification (Monarch® Plasmid Miniprep Kit, New England Biolabs, #T1010) and sequencing (Source BioScience PLC). Quantitative real time PCR (qPCR) was used to confirm reduction of *rbfox1* mRNA expression.

### Fish breeding and genotyping

For adult experiments, mixed sexes were used. When possible, animals were associated with an identification number and genotype was assigned after data analysis. For larval experiments, fish were generated by heterozygous in-cross and genotyped prior to data analysis. Genomic DNA was extracted from fins and using the HotSHOT method. Briefly, samples were incubated at 95 °C in 50 mM NaOH for 30 min, followed by 1 min at 10 °C. Reaction was stopped using 1M Tris HCl (1/10 of the initial NaOH volume), pH 8.00. Genotyping primers were the same as the ones used to identify founder carriers.

### Drug Treatment

Zebrafish embryos of each genotype (*rbfox1*^*+/+*^ and *rbfox1*^*19del/19del*^) were treated from 5hpf to 5dpf with 20 μM TRKB antagonist ANA-12 (abcam, Cat.ab146196), or 20 μM TRKB agonist 7,8-dihydroxyflavone (7,8-DHF) (abcam #ab120996), or dimethylsulfoxide (DMSO, vehicle 0.01%) (Merck #34869). Treatment was performed in Petri dishes (max 30 embryos x dish) and drugs or vehicle were dissolved in fish water. Solutions were replaced daily.

### Quantitative Real-Time PCR

Quantitative Real-time PCR (qPCR) of target RNA was performed on 5dpf *rbfox1* wild type (*rbfox1*^*+/+*^) and rbfox1 homozygous (*rbfox1*^*19del/19del*^) larvae, using Luna® Universal One-Step RT-qPCR Kit (New England Biolabs #M3005) and a Bio-Rad 96-well qPCR machine (CFX96 Touch Real-Time PCR Detection System). Total RNA was isolated using TRIzol reagent (Thermo Fisher Scientific) following manufacturer’s instructions. Briefly, after homogenization, RNA was isolated by precipitation, rinsed in ethanol and resuspended in RNase free water. Total RNA was then quantified using BioDrop (Biochrom Ltd.), and up to 1 μg was reverse transcribed to cDNA using the ProtoScript II First Strand cDNA Synthesis Kit (New England Biolabs, #E6560) following manufacturer’s instructions. The resulting cDNA yield and quality were evaluated using BioDrop (Biochrom Ltd.). As for zebrafish trkb2 *TK+* and *TK-* isoforms quantification, Ct values resulting from the amplification of the *TK+* specific product were subtracted from the Ct values resulting from the amplification of the *TK+*/*TK-* common product. All reactions included 5 biological replicates and 3 technical replicates. For experiments in larvae, each biological replicate consisted of 15 larval heads (eyes and jaw removed). For experiments in adults, each biological replicate consisted of a single brain. Actin – β 2 (*actb2*) and ribosomal protein L13a (*rpl13a*) were employed as reference genes. For assessing *rbfox1* mRNA expression, *rbfox1* primers were designed upstream of the CRISPR deletion. Accession numbers, primer sequences and amplification efficiencies for all the reference and target genes can be found in Supplementary Table 1.

### In situ hybridization

*In situ* hybridization (ISH) was carried out on whole mount zebrafish larvae and on larval (sagittal) and adult brain (transverse) sections as described previously2. For *rbfox1* ISH, the original plasmid used to generate the riboprobe was provided by Dr William Norton (University of Leicester). The plasmid used to produce the *trkb2* riboprobe was generated in our laboratory (Forward primer: 5’-GTTCGTGGAATGGCTTGCTG-3’, Reverse primer: 5’-TCTGGCCCACGATGTTTTCA-3’) using the pGEM®-T Easy Vector System (Promega, #A1360) and *in-house* generated *E. coli* DH5α competent cells. Riboprobes to identify *rbfox1* (NM_001005596) and *trkb2* (NM 01197161.2) mRNA were synthetized by in vitro transcription (IVT) using MAXIscript™ SP6/T7 kit (Invitrogen by Thermo Fisher Scientific, #AM1322), following manufacturer’s instructions and using a DIG RNA Labeling Mix, 10× conc (Roche, #11277073910) containing digoxigenin labeled uracil.

### In situ hybridization on whole mount zebrafish larvae

*In situ* hybridization was carried out on 28hpf, 2-3-4- and 5dpf *rbfox1*^*+/+*^ and *rbfox1*^*19del/19del*^ larvae. To prevent skin pigmentation, embryos were incubated in 0.2 mM 1-phenyl 2-thiourea (PTU) (Sigma, #S527335) from 24hpf. When they reached the desired age, larvae were fixed in 4% paraformaldehyde (PFA) (Merck, #158127) overnight (ON) at 4°C. The following day, larvae were rinsed in 1x phosphate buffered saline (PBS) (Thermo Fisher Scientific, #18912014) supplemented with Tween 20 (Sigma, #P1379) (0.05% v/v), dehydrated in ascending methanol series (25%, 50%, 70%, 80%, 90%, 100% methanol, 5 min each) and stored in 100% methanol at -20 °C. To perform ISH experiments, larvae were rehydrated in descending methanol series (100%, 90%, 80%, 70%, 50%, 25% methanol), 5 min each, and washed in 1xPBS, 5 min. Larvae were permeabilized using proteinase K (PK) (ITW Reagents, #A3830) (stock 20 μg/mL in 1xPBS) as follows: 28hpf larvae were permeabilized in PK 1:2000 in 1xPBS for 20 min at room temperature (RT), older stages were permeabilized in PK 1:1000 at 37°C for at least 30 min. Then, larvae were post fixed in 4% PFA for 20 min and washed in 1xPBS at RT, 5 × 5 min. Prehybridization was carried out in hybridization solution (HB) containing 50% formamide, 5% saline sodium citrate buffer (SSC), 50 µg/mL heparin, 0.05 mg/mL yeast RNA, 0.1% Tween 20, and 0.92% citric acid at 68 °C for 2h. Thereafter, larvae were incubated in HB containing *rbfox1* riboprobe (500 pg/µL), ON at 68°C. Post hybridization washes were performed at 68°C with a gradient of 2xSSC and formamide (50%, 25% and 0% formamide), 10 min each, and then twice with 0.02xSSC, 30 min each. Subsequently, larvae were blocked in blocking solution (BS) containing 10% normal sheep serum (Gibco, #16070096) and 2 µg/µL bovine serum albumin, for 1h at RT. After blocking step, larvae were incubated in anti-digoxigenin Fab fragments conjugated with alkaline phosphatase (Roche, #11093274910), 1:2000 in BS, 1h at RT and then ON at 4°C. The following day, larvae were washed in 1xPBS, 6 × 15 min each, and then in alkaline phosphatase (100 mM NaCl, 100 mM Tris HCl, 50 mM MgCl2, 0.1% Tween 20) (NTMT) buffer, 3 × 5 min each. The chromogenic reaction was carried out by incubating the larvae in BCIP/NBT solution (Merck, #203790) in NTMT buffer, at RT in the dark, and were observed every 20 min until the signal detection. After reaching the desired staining, larvae were washed in 1xPBS at RT, post fixed in 4% PFA for 2h, cleared and stored in 80% glycerol at 4°C. For sagittal cryosections, larvae were embedded in 1.5% low gelling temperature agarose (Scientific Laboratories Supplies, #A9414) supplemented with 5% Sucrose in 1xPBS. Sections (5 μm) were collected on adhesive microscope slides Superfrost® Plus Gold (Epredia).

### In situ hybridization on sections

ISH was conducted on paraffin embedded *rbfox1*^*+/+*^ and *rbfox1*^*19del/19del*^ adult brains (transverse sections) and 5dpf *rbfox1*^*+/+*^ and *rbfox1*^*19del/19del*^ zebrafish larvae (sagittal sections). Fish were culled by an overdose of tricaine prior to head removal. Dissected adult brains and larvae were fixed in 4% PFA in 1xPBS, ON at 4°C. Tissues were then rinsed in 1xPBS and dehydrated in ascending ethanol series (15 min in each of 30%, 50%, 70%, 80%, 90%, 100% ethanol) and embedded in paraffin. Transverse (adult brains) or sagittal (5 days old larvae) sections of 12 µm (adult brains) or 7 µm (5 days old larvae) thickness were cut using a microtome (Leica). To perform ISH, slides were de-waxed in xylene (twice, 10 min each), rehydrated in descending ethanol series (2 × 5 min in absolute ethanol, then 90%, 80%, 70%, 50% and 25% ethanol, 5 min each), and rinsed in 1xPBS for 5 min. Then, sections were permeabilized using PK (0.05 μg/μL) for 8 min at RT, washed with 2 mg/mL glycine twice (5 min each), post fixed in 4% PFA for 20 min and washed in 1xPBS at RT. Prehybridization was carried out in HB, for 1h at 68°C. Thereafter, sections were incubated in HB containing *rbfox1* riboprobe (500 pg/µL), ON at 68°C. Post hybridization washes were performed at 68°C twice for 20 min in 1xSSC, twice for 20 min in 0.2xSSC, and several washes were performed in 1xPBS, 5 min each at RT. Then, sections were blocked in BS for 30 min at RT and incubated in a 1:2000 dilution of anti-digoxigenin Fab fragments conjugated with alkaline phosphatase in BS, ON at 4°C. The following day, sections were washed in 1xPBS, 5 × 10 min each. The chromogenic reaction was carried out by incubating the slides in BCIP/ NBT solution in NTMT buffer, at RT in the dark, and were observed every 20 min until the signal detection. When the desired staining was obtained, sections were washed in 1xPBS at RT, dehydrated in ascending ethanol series (25%, 50%, 70%, 80%, 90%, 100% ethanol, 5 min each), cleared in xylene (twice, 5 min each) and mounted with dibutyl phthalate polystyrene xylene mounting medium (Sigma, #06522).

### Image acquisition and processing

Pictures of whole mount ISH on zebrafish larvae were acquired by Leica MZ75 microscope. For ISH on sections, pictures were acquired using a Leica DMRA2 upright epifluorescent microscope with colour QIClick camera (Leica) and processed with Velocity 6.3.1 software (Quorum Technologies Inc). Quantification of the ISH staining signal intensity was performed as described previously72, using Fiji software73. Adult anatomical structures were identified according to the Neuroanatomy of the Zebrafish Brain by Wullimann 74.

### Behavioral Assays

For larvae, all behavioural experiments were conducted on the progeny of a *rbfox1*^*+/19del*^ in-cross. Larvae were genotyped prior to data analysis. For adults, where the genotype of the animals was known, fish were pseudorandomised across testing systems with all trials having an approx. equal number of each genotype. The adult fish were weight- and age-matched, with approximately equal numbers of both sexes. Sex was not treated as a biological variable in the statistical analyses.

### Larval behavioural experiments

Patterns of locomotor activity of 5dpf *rbfox1*^*+/+*^, *rbfox1*^*+/19del*^ and *rbfox1*^*19del/19del*^ mutant zebrafish larvae were investigated as described previously24,29,33. Tests were conducted between 9 a.m. and 4 p.m. At 5 dpf, larvae were placed in individual wells of a 24-well plate. To reduce stress due to manipulation, larvae were acclimatised for at least 1 h before testing. Then, plates were placed into the DanioVision observation chamber (Noldus). Locomotion parameters such as distance travelled and swimming velocity were recorded using EthoVision XT software (Noldus). Data were exported in 1 min and 1 sec time bins and analysed with R programming language75. For the hyperactivity assay, larval basal locomotion was tracked for 15 min in dark conditions. Larval swimming burst was assessed as described previously24 and peaks were considered as the acceleration events when larvae travelled > 5 mm in < 12 sec. The forced light-dark transition assay was performed as described previously76,77 with modifications: after an initial 5 min period of dark (baseline), larvae were exposed to one light/dark cycle of 1 min light (Noldus white light device) followed by 5 min dark. The response and habituation to acoustic startle stimuli was performed as described previously29: after 10 min of baseline (no stimuli, dark conditions), larvae were subjected to 10 sound/vibration stimuli (Noldus tapping device) over 20 sec (2 sec intervals between each stimulus).

### 5-Choice Serial Reaction Time Task

We measured impulsive action using a zebrafish version of the 5-Choice Serial Reaction Time Task (5-CSRTT)25. Adult *rbfox1*^*+/*+^, *rbfox1*^*+/19del*^ and *rbfox1*^*19del/19del*^ zebrafish, 7 months old, mixed sexes, were singly housed for a week prior to experiment and remained singly housed for the whole duration of the assay (9 weeks). Fish were tested using the Zantiks AD units. Each unit was provided with a small tank with five apertures and a food hopper insert. The five apertures created five different entry points for the fish, acting like the five nose poke holes of the rodent version of the assay. The food hopper was placed at the opposite side of the five apertures and formed an area for the fish to enter and collect food reward. Below the testing tank there was an integrated screen, used to display white light (stimulus) into the five apertures. Responses were detected when a fish entered these apertures and recorded with an integrated camera placed at the top of the tank. The experiment consisted of five training stages: i) habituation, ii) initiator training, iii) stimulus light training, iv) 5-CSRTT/no delay, v) 5-CSRTT/variable-delay. Details of each stage are provided in Supplementary Table 2.

### Novel tank diving

Novel tank diving is a behavioural test to assess anxiety-like behaviour in adult fish. Response to novel tank was assessed in *rbfox1*^*+/+*^, *rbfox1*^*+/19del*^ and *rbfox1*^*19del/19del*^ 9 months old zebrafish, mixed-sexes, as described previously33. Fish were singly housed for a week prior to performing the experiment and acclimatized for at least 1 h in the behavioral room on the testing day. Behavioural assays were conducted between 9 a.m. and 2 p.m. During the test, fish were individually placed into a 1.5 L tank and their behaviour was tracked and recorded using EthoVision system (Noldus). Data were exported in 1 min time bin and analysed as previously described33. Experimental groups were randomised during testing. We analyzed three behaviours in response to the novel tank: i) time spent in the bottom of the tank, ii) total distance traveled, and iii) number of transitions to the top– bottom area of the tank.

### Whole-body cortisol extraction

#### Larvae

Larval cortisol was extracted from pools of 12 larvae x sample using a modified protocol based on Baiamonte et al. (2016)35. Briefly, larvae were snap-frozen upon collection and stored at -20⍰°C until extraction was performed. On the day of extraction, frozen samples were thawed on ice in and homogenised in 400⍰μl of ice-cold 1×PBS for 30 seconds using a Bioruptor (MP Biomedicals). Cortisol was extracted by adding 500⍰μl of ethyl acetate (Sigma) to each homogenate. Samples were vortexed for 30 seconds, centrifuged at 5000⍰rpm for 10 minutes and then snap frozen at −80⍰°C. The organic phase was carefully decanted into glass vials (VWR). The extraction was repeated twice and the combined organic phases were pooled. The pooled ethyl acetate extracts were evaporated at 60⍰°C using a rotary evaporator (IKA). The dried cortisol was resuspended in 200⍰μl of 1xPBS, vortexed for 30 seconds and stored at −20⍰°C until cortisol quantification using a human salivary cortisol ELISA kit (Salimetrics).

#### Adults

Adult cortisol was extracted from 4 adult zebrafish whole-body. Fish were weighed and homogenised in 2× volumes (w/v) of 1xPBS using a Bioruptor (MP Biomedicals). Cortisol was extracted using the same ethyl acetate procedure as for larvae. However, after evaporation, to prevent any interference with the assay, residual lipids were eliminated by partitioning the dried extracts between 500μl 1xPBS and 500μl hexane (Sigma). The upper organic layer was discarded and the aqueous phase was retained and stored at −20⍰°C until cortisol quantification using a human salivary cortisol ELISA kit (Salimetrics).

### Cortisol measurement

Samples were thawed on ice and 50⍰μl of each was assayed using the Salimetrics human salivary cortisol ELISA kit, following the manufacturer’s protocol. Cortisol values for adults were normalised to body weight (ng/g), while larval and juvenile values were normalised to total tissue protein (ng/mg). This assay was validated for use with zebrafish whole-body extracts previously35,78. Standards, high and low controls and zebrafish samples (25⍰μL) were added to the plate provided with the Salimetric kit, along with Assay Diluent (25⍰μL) for the zero standard. The Enzyme Conjugate was diluted 1:1600 in Assay Diluent and 200⍰μL was dispensed into each well. After mixing for 5⍰minutes at 500⍰rpm, the plate was incubated at RT for 1h and washed four times with 1× wash buffer (300⍰μL x well x wash). Next, 200⍰μL of TMB Substrate Solution was added, followed by a 30-minute dark incubation with first 5min mixing. The reaction was stopped with 50⍰μL of Stop Solution, mixed until wells turned yellow, and absorbance was read at 450⍰nm within 10⍰minutes using an OMEGA plate reader (BMG Labtech).

### Statistical analysis

For qPCR, relative mRNA expressions were calculated using the Pfaffl method79. Outliers were identified and removed using Dixon’s test (α = 0.05)80. Differences in gene expression were assessed using a one-way ANOVA followed by Tukey’s post-hoc test using GraphPad Software (Prism). For behavioural analysis, all data were analysed with R programming language75. Scripts used for analysis are available on GitHub (https://github.com/AdeleLeg). For models where distance moved, distance to zone, velocity, or top halves visits were the response variable, we fitted data to mixed linear models using the “*lme4*” package, and where proportion of responders or proportion of time spent in the bottom third were our response variable, we fitted data to beta regression models using the “*betareg*” package. In all instances, for all experiments, we used genotype as fixed effect, and fish ID and day of experiment as random effects. In the response and habituation to acoustic startle, we used also the stimulus number as fixed effect. As in García-González et al.33, we reported significant fixed effects as Type II Wald χ2 from models using the package “*car*,” post hoc Tukey’s tests were also conducted as necessary with the package “*emmeans*”. For the 5-CSRTT, overall correct responses (learning) and anticipatory responses (impulsivity) were assessed using the formulas in Supplementary Table 2. Statistical analysis of mean intensity in in situ hybridization images of zebrafish larval sections was performed using ImageJ. Mean intensity values were quantified across specified regions of interest, indicated in Supplementary Figure 5, ensuring that comparable areas of similar dimensions were used for each sample to calculate differences in expression. The resulting values were compared across groups using a paired t-test to determine significant differences using GraphPad Software (Prism). For all experiments, sample size needed to achieve adequate statistical power for detecting a significant effect was determined based on data from previous research or pilot studies. Accepted α level and power were respectively ≤ 0.05 and ≥ 0.80.

### Fertilisation and survival analysis

Fertilisation and survival rates were measured as described previously19. For fertilization rate we measured the percentage of fertilized eggs when pairing *rbfox1*^*19del/19del*^ and compared it with fertilization rate of *rbfox1*^*+/+*^ siblings. Data were collected upon 3 different mating trials, each trial comprising 3-5 trios (1 male and 2 females). Trials were performed on different weeks, using different trios for each test. Eggs were collected in different Petri dishes, properly labelled to distinguish between trios. Fertilisation rate was determined by averaging across Petri dishes from the same trio and calculated at 6-8hpf. For survival analysis, we measured the percentage of surviving animals (across 3 tanks for each genotype for each batch, to avoid tank effect) from when larvae were added to the nursery (at 5dpf) until 2 months of age, when fish were transferred into adult aquarium.

## CONTRIBUTIONS

AL conceived the project, designed and conducted the experiments, analysed the data, and wrote the manuscript. JGG designed and performed the CRISPR/Cas9 experiment generating the line. SH, PA, SA, XW and WH contributed to the experiments. NFC and BC edited the manuscript. CHB directed the study, designed the experiments, edited the manuscript, and secured funding. All authors contributed to the article and approved the submitted version.

### ACKNOWLEDGMENTS

CHB, AL, SH, PA and WH were supported by the NIH grant U01 DA044400-03. JGG was supported by a 2022 NARSAD Young Investigator Grant (Number #30749) by the Brain & Behavior Research Foundation. NFC and BC were supported by the grants PID2021-127776OB-I100, 202218 31 and 2021-SGR-01093 from the Ministry of Science, Innovation and Universities (Spain), Fundació La Marató de TV3, and AGAUR-Generalitat de Catalunya, respectively, and by ICREA Academia 2021. Authors would like to acknowledge SciDraw (https://scidraw.io/) for images used to create the graphical abstract. We thank Prof. Marina Resmini for kindly providing the equipment used to dry cortisol extracts.

## CONFLICT OF INTEREST

The authors declare that the research was conducted in the absence of any commercial or financial relationships that could be construed as a potential conflict of interest.

---

**Supplementary Figure 1.**
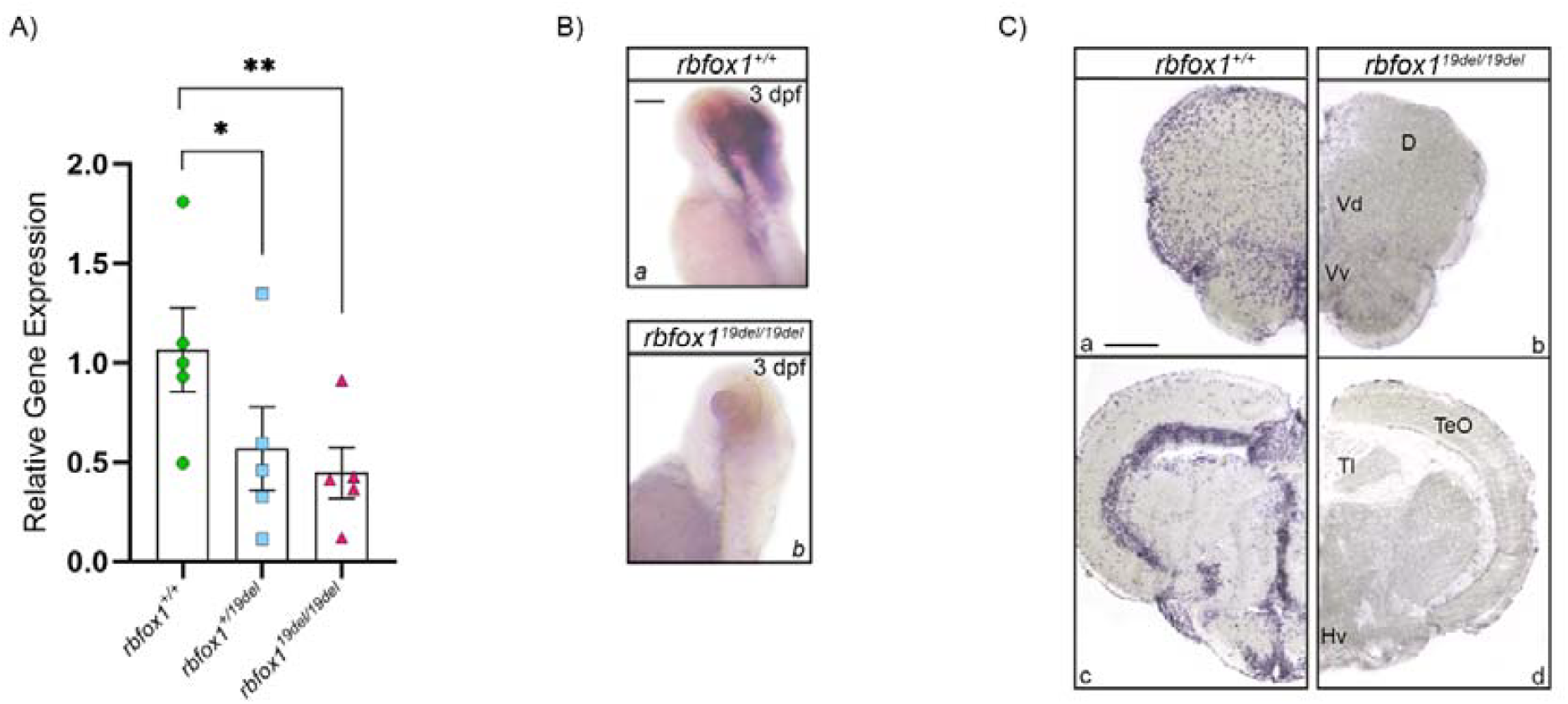
Generation of a *rbfox1* loss of function line. **A)** *rbfox1* mRNA expression in *rbfox1*^+/+^, rbfox1+/19del and rbfox1^19del/19del^ 5 days post fertilisation (dpf) larvae. In *situ* hybridisation on **B)** whole mount zebrafish larvae at 3dpf, a) *rbfox1+/+* and b) *rbfox1*^19del/19del^ and **C)** adult transverse sections of *rbfox1+/+* (a) fore- and (c) mid-brain and *rbfox1*^19del/19del^ (b) fore- and (d) mid-brain. In A) each dot/square/triangle represents a pool of *rbfox1+/+* or *rbfox1*^19del/19del^ 15 larval heads (eyes and jaw removed) respectively. For qPCR experiments, reference genes were actin – β 2 (*actb2*) and ribosomal protein L13a (*rpl13a*). All larvae employed were progeny of *rbfox1+/19del* in-cross and were genotyped after experiments and prior to data analysis. In A): bars represent standard error of the mean (SEM); * p < 0.05; ** p < 0.01. Scale bars: 200 µm in B); 100 µm in C). Abbreviations: D, dorsal area of dorsal telencephalon; Hv, ventral hypothalamus; TeO, optic tectum; Tl, longitudinal tori; Vd, ventral area of dorsal telencephalon; Vv, ventral area of ventral telencephalon. N = 4 x genotype.

**Supplementary Figure 2.**
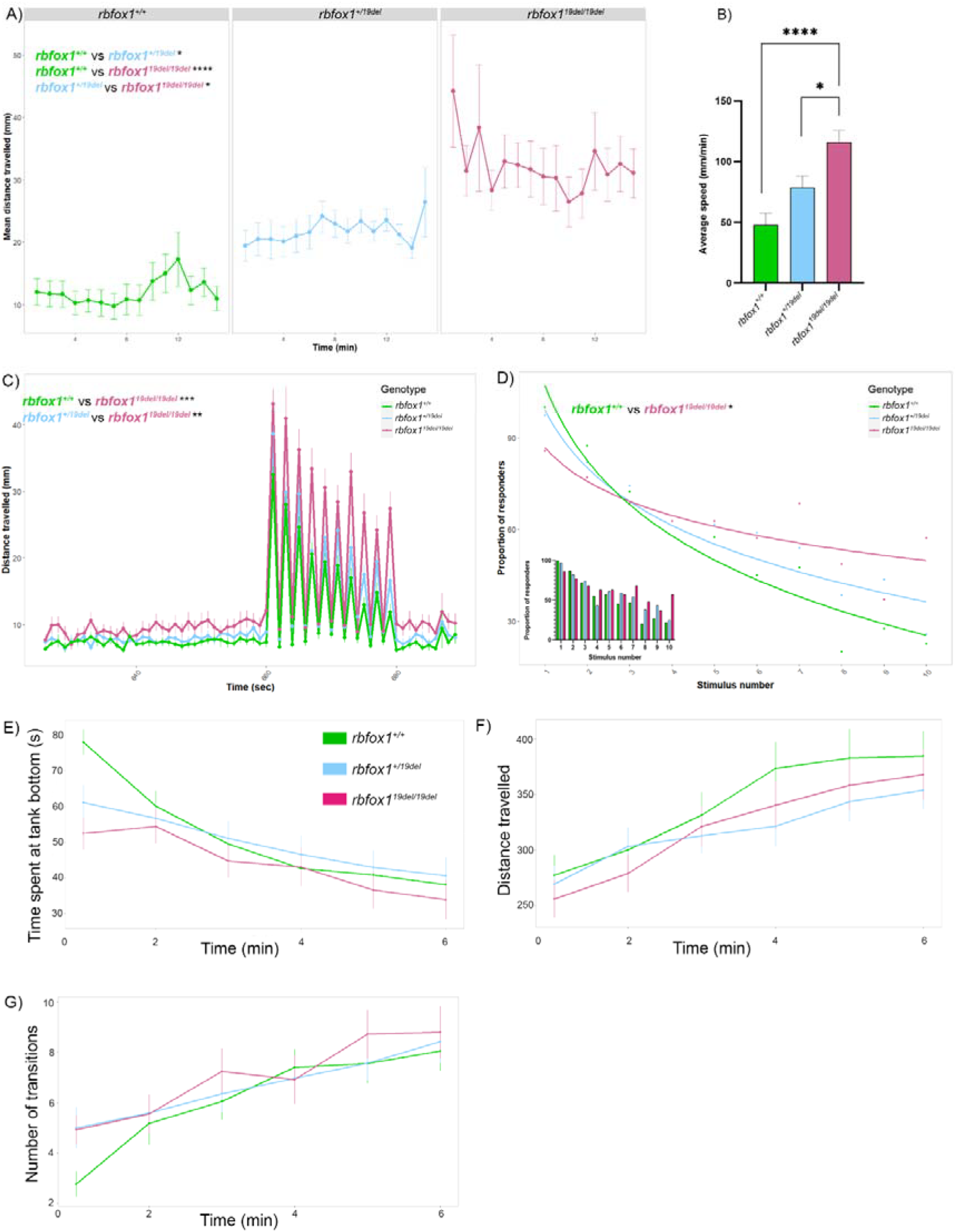
rbfox1 mutant fish show hyperactivity, impulsivity and hyperarousal behaviour. **A-B)** Locomotion assay in 5 days post fertilization (dpf) zebrafish larvae (*rbfox1*^*+/+*^, *rbfox1*^*+/19del*^, *rbfox1*^*19del/19del*^): **A)** distance travelled during 15 min; **B)** average speed; N (A-B) = *rbfox1*^*+/+*^ *24; rbfox1*^*+/19del*^ 24; *rbfox1*^*19del/19del*^ 24 5-9). **C-D)** Response and habituation to acoustic startle assay in 5dpf zebrafish larvae (*rbfox1*^*+/+*^, *rbfox1*^*+/19del*^, *rbfox1*^*19del/19del*^): **C)** mean distance travelled during the assay; **D)** rate of habituation over time/stimuli; N = *rbfox1*^*+/+*^ 40; *rbfox1*^*+/19del*^ 39; *rbfox1*^*19del/19del*^ 35. E-G) Novel tank diving assay in *rbfox1*^*+/+*^, *rbfox1*^*+/19del*^ and *rbfox1*^*19del/19del*^. **E)** Time spent at the bottom of the tank. **F)** Distance travelled during the assay and **G)** number of transitions between the top and bottom of the tank. N = *rbfox1*^*+/+*^, 50; *rbfox1*^*+/19del*^ 54; *rbfox1*^*19del/19del*^ 54. Bars represent standar error of the mean (SEM). All larvae employed in behavioural experiments were progeny of *rbfox1*^*+/19del*^ in-cross and were genotyped after experiments and prior to data analysis. In all graphs: bars represent standard error of the mean (SEM); * p < 0.05; ** p < 0.01; *** p < 0.001; **** p < 0.0001.

**Supplementary Figure 3.**
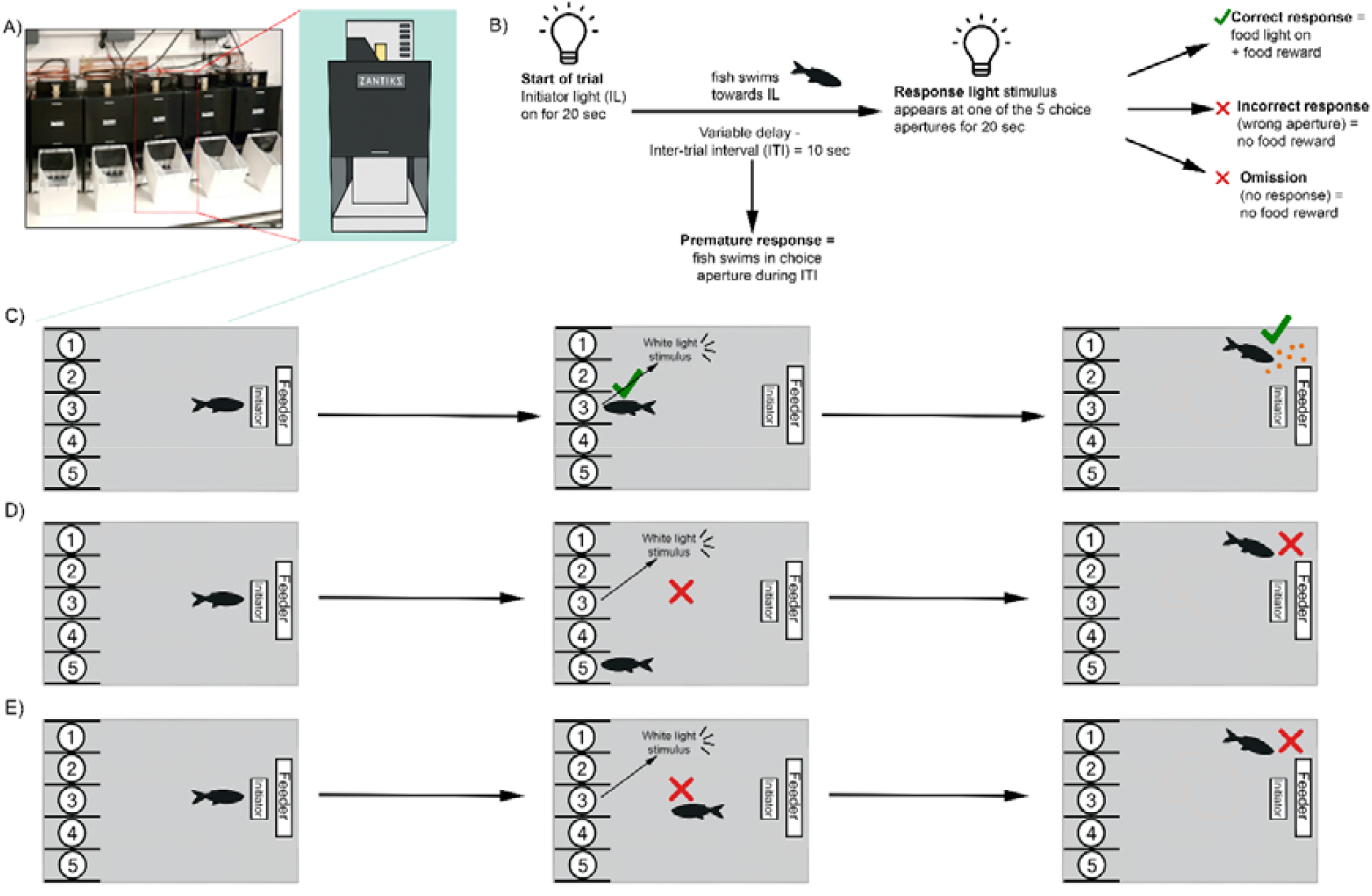
5-choice serial reaction time task. **A)** Zantiks behavioural testing apparatus used in the laboratory (left panel) and a schematic representation of a single Zantiks chamber (right panel). **B)** Schematic overview of the pipeline in a single 5-choice trial, including initiation, stimulus presentation and response outcome. **C-E)** Illustrative diagrams of representative trial outcomes as depicted in the pipeline in panel B), including C) correct response, D) incorrect response and E) omission.

**Supplementary Figure 4.**
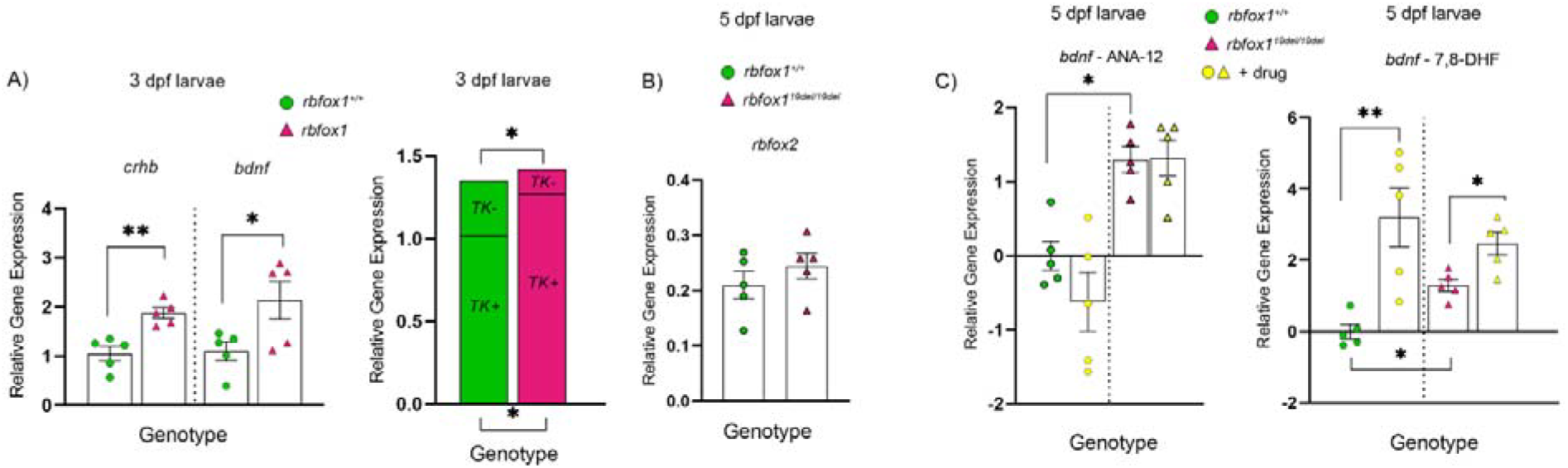
**A)** *crhb* and *bdnf* (left) and trkb2 short and long isoform (right) expression levels in 3 days old larvae, *rbfox1+/+* and *rbfox1*^*19del/19del*^. **B)** *rbfox2* expression levels in 5 days old larvae, *rbfox1+/+* and rbfox1^*19del/19del*^. **C)** *bdnf* expression levels in 5 days old larvae, *rbfox1+/+* and rbfox1^*19del/19del*^, following chronic exposure to either ANA-12 (left) or 7,8-DHF (right). Actin – β 2 (*actb2*) and ribosomal protein L13a (*rpl13a*) were used as reference genes. Dots/triangles represent pool of 15 larval heads (eyes and jaw removed). Bars represent standard error of the mean (SEM). N = *rbfox1+/+* 5; *rbfox1*^*19del/19del*^ 5.

**Supplementary Figure 5.**
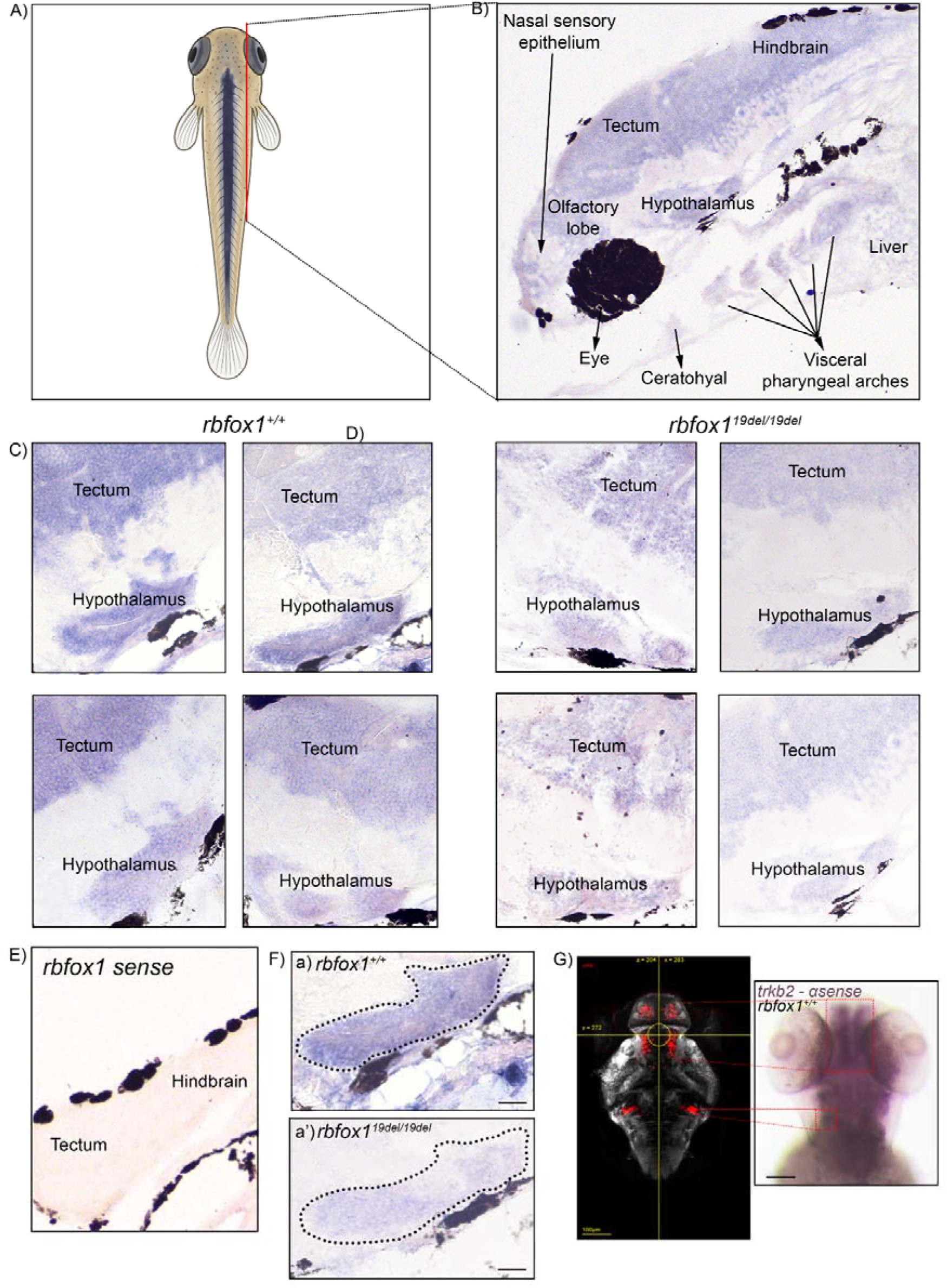
Details of *in situ* hybridisation experiments for *trkb2* anti-sense riboprobe on 5 days post fertilisation zebrafish larvae, sagittal sections. **A)** Schematic representation indicating the level of the anatomical section (red line). **B)** Representative sagittal section of zebrafish larva stained for *rbfox1* (anti-sense riboprobe), with anatomical regions labelled in black according to ZFIN atlas of zebrafish larvae (https://zfin.org/zf_info/anatomy.html). **C-D)** Overview of zebrafish larvae hypothalamus (sagittal sections), *rbfox1*^*+/+*^ (four panels on the left) and *rbfox1*^*19del/19del*^ (four panels on the right). Each panel corresponds to a different animal. N = 4 per genotype. Sections have been placed on the same slide and were stained under the same conditions. **E)** *In situ* hybridisation for *trkb2* sense riboprobe on 5 days post fertilisation zebrafish larva, sagittal section. **F)** Example of areas selected (areas enclosed by the black dotted lines) to calculate *trkb2* mean intensity in the hypothalamus of a) zebrafish *rbfox1*^*+/+*^ (left panel) and a’) *rbfox1*^*19del/19del*^ (right panel) larval sections. **G)** Comparison of *trkb2* and *crhb* expression in the larval zebrafish brain. The left panel displays *crhb*-expressing cells in the larval brain from the publicly available MapZebrain atlas (https://mapzebrain.org/atlas/2d). The right panel shows whole-mount *in situ* hybridisation for *trkb2* expression in 5 days old zebrafish larva. Scale bars: F) 50 µm; G) 100 µm.

**Supplementary Table 1.**
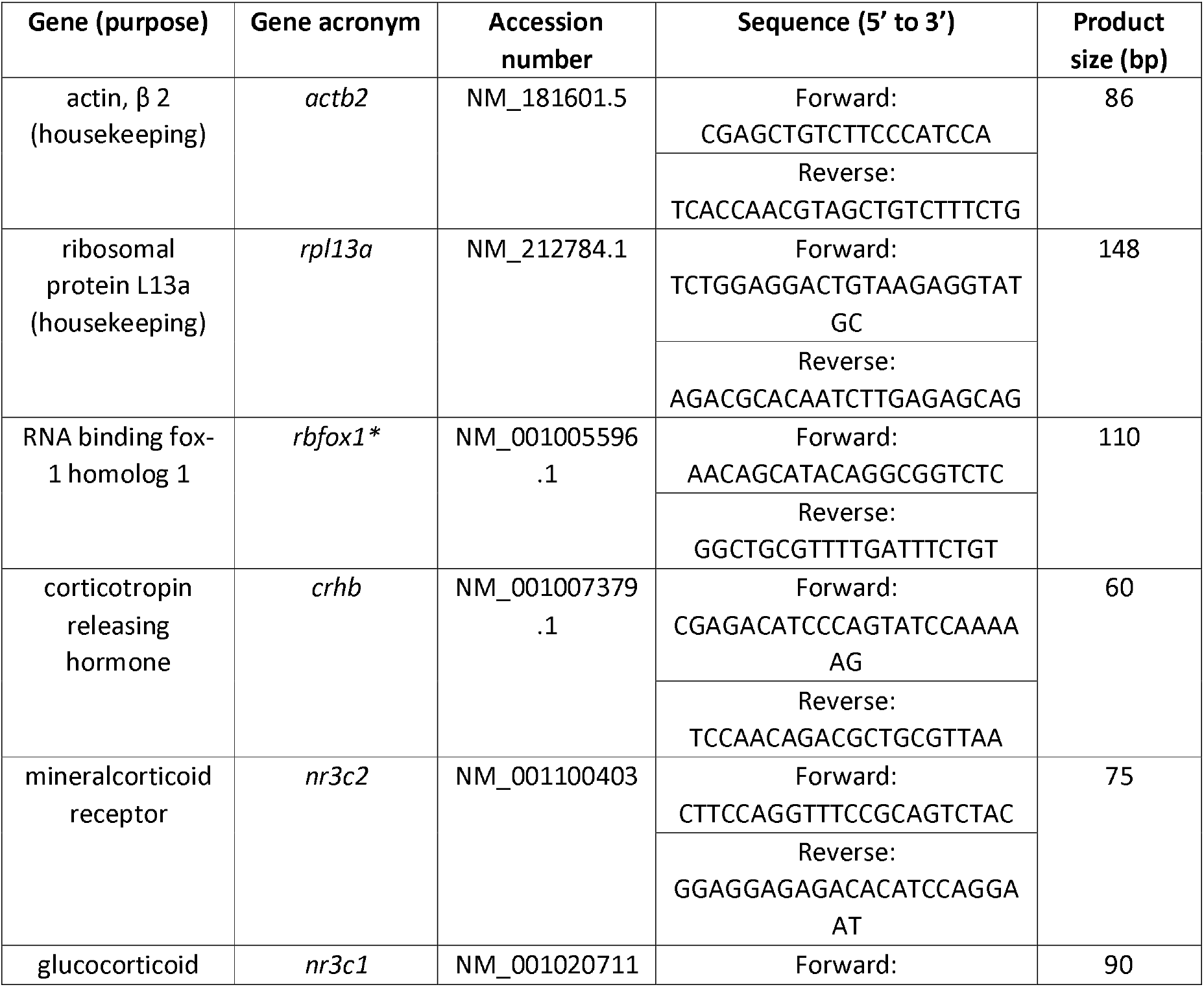
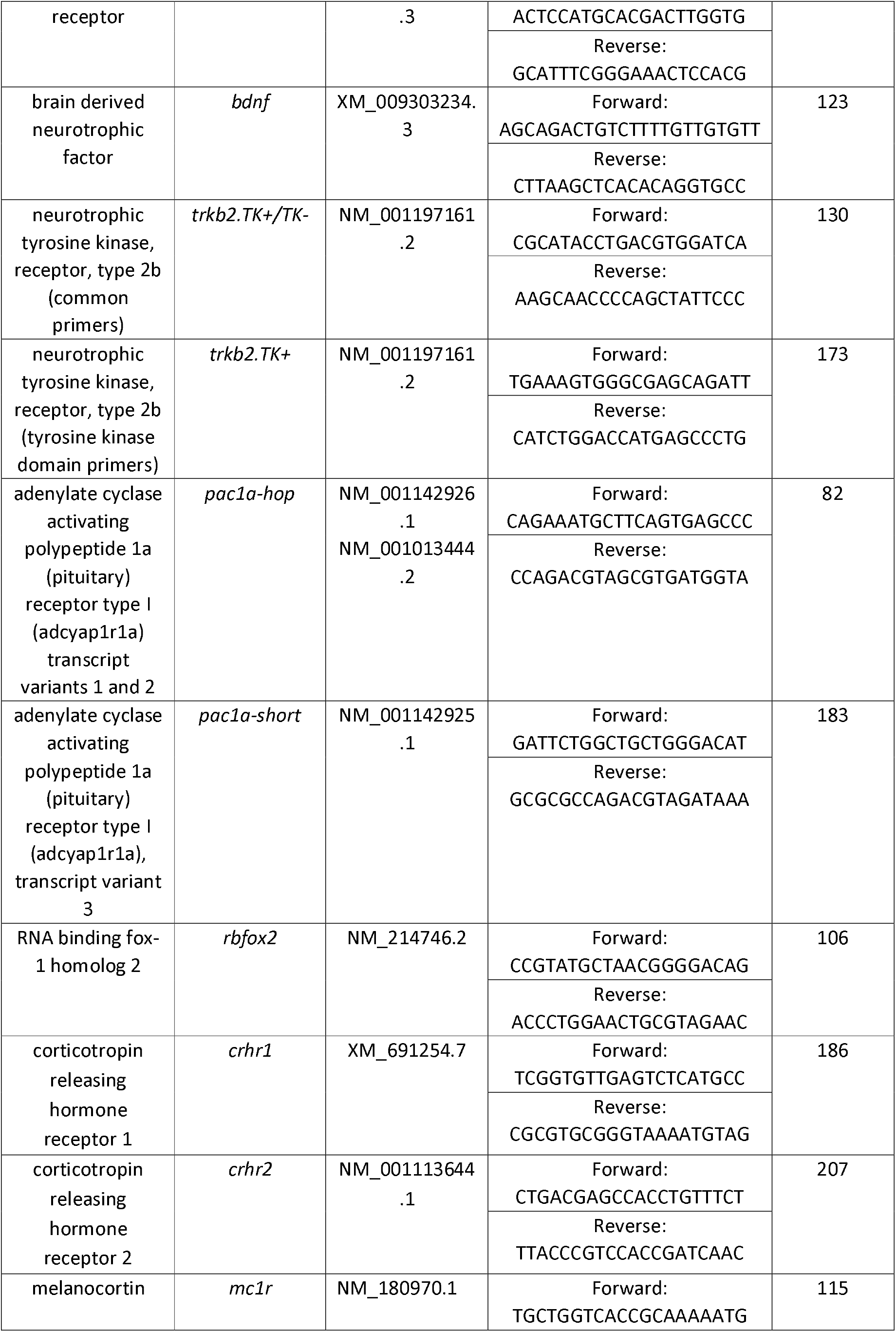
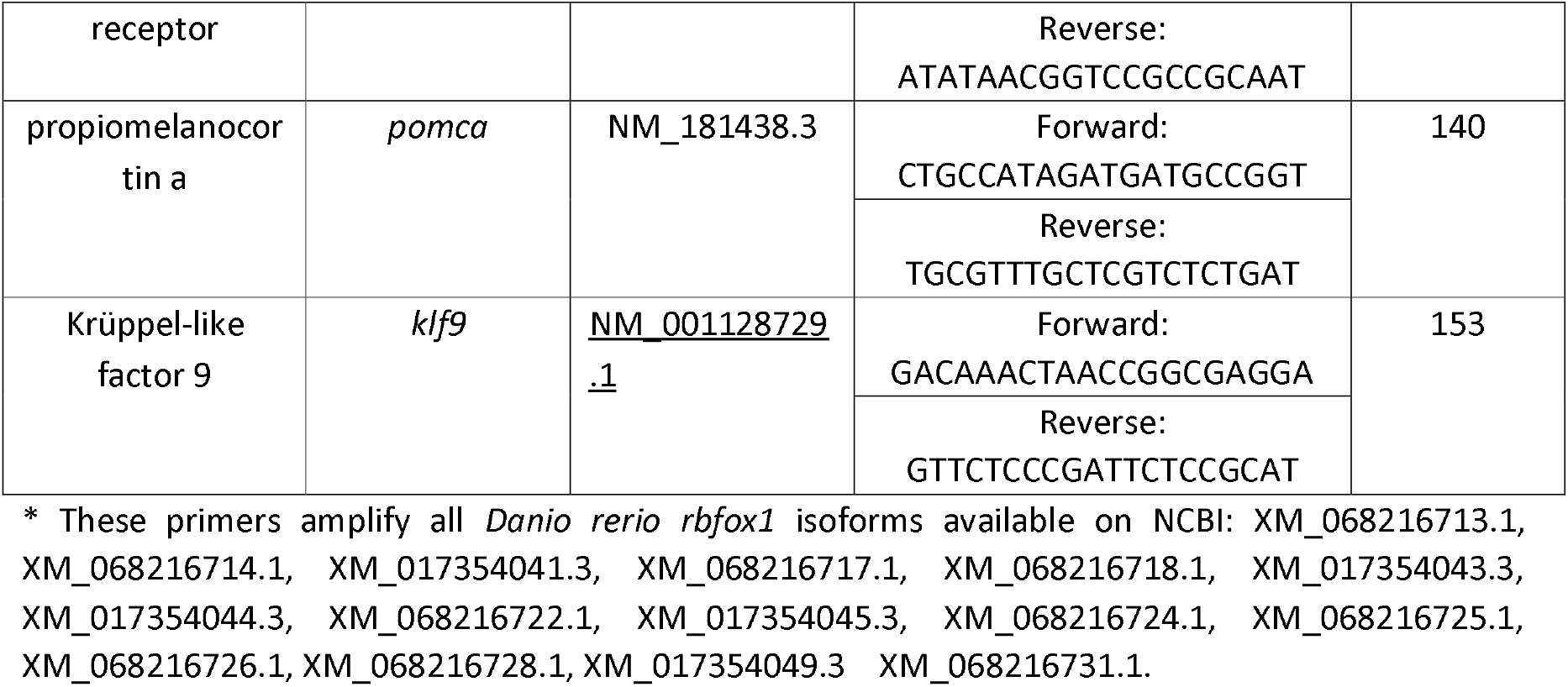
List of primers used for qPCR experiments.

**Supplementary Table 2.**
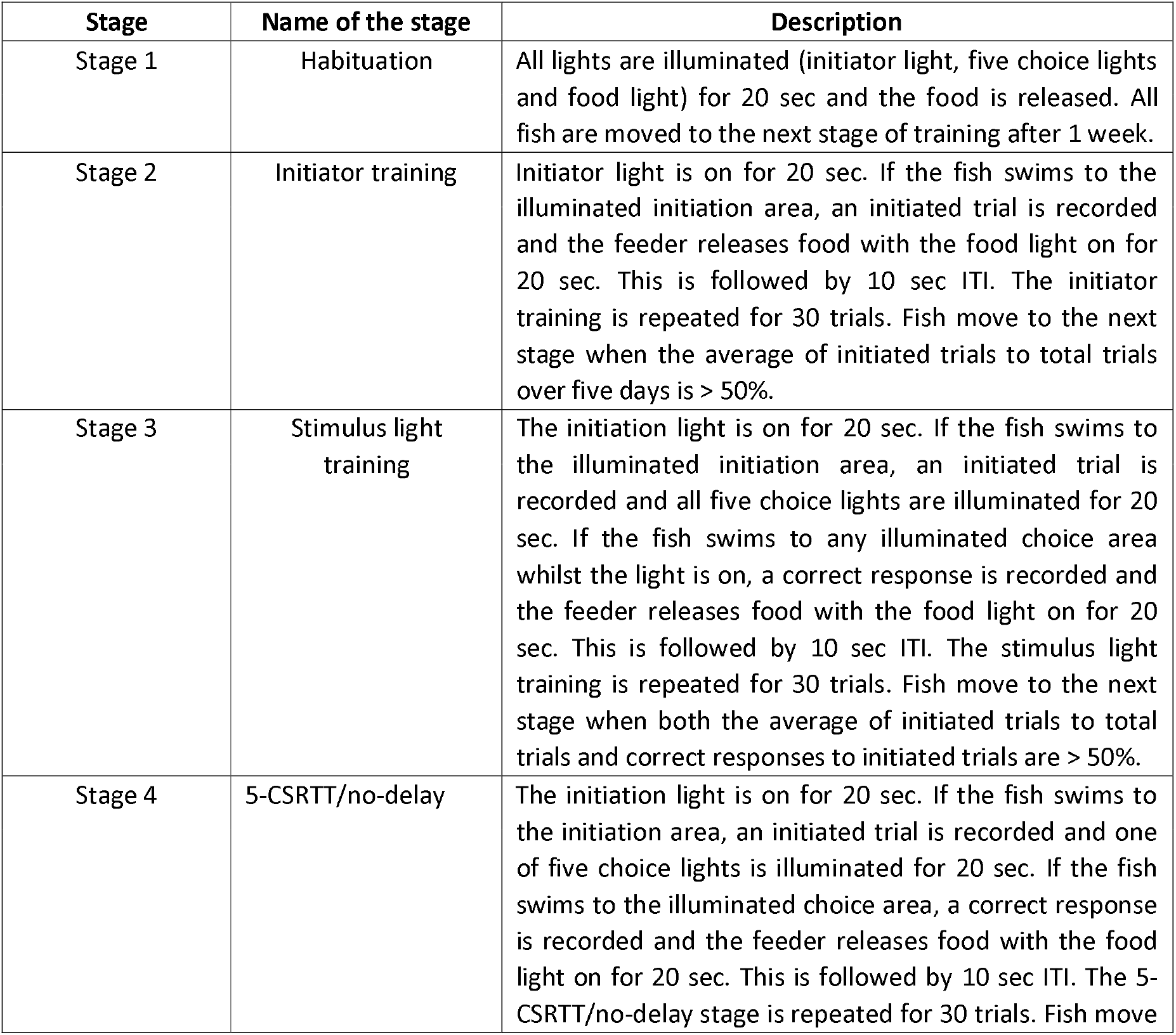
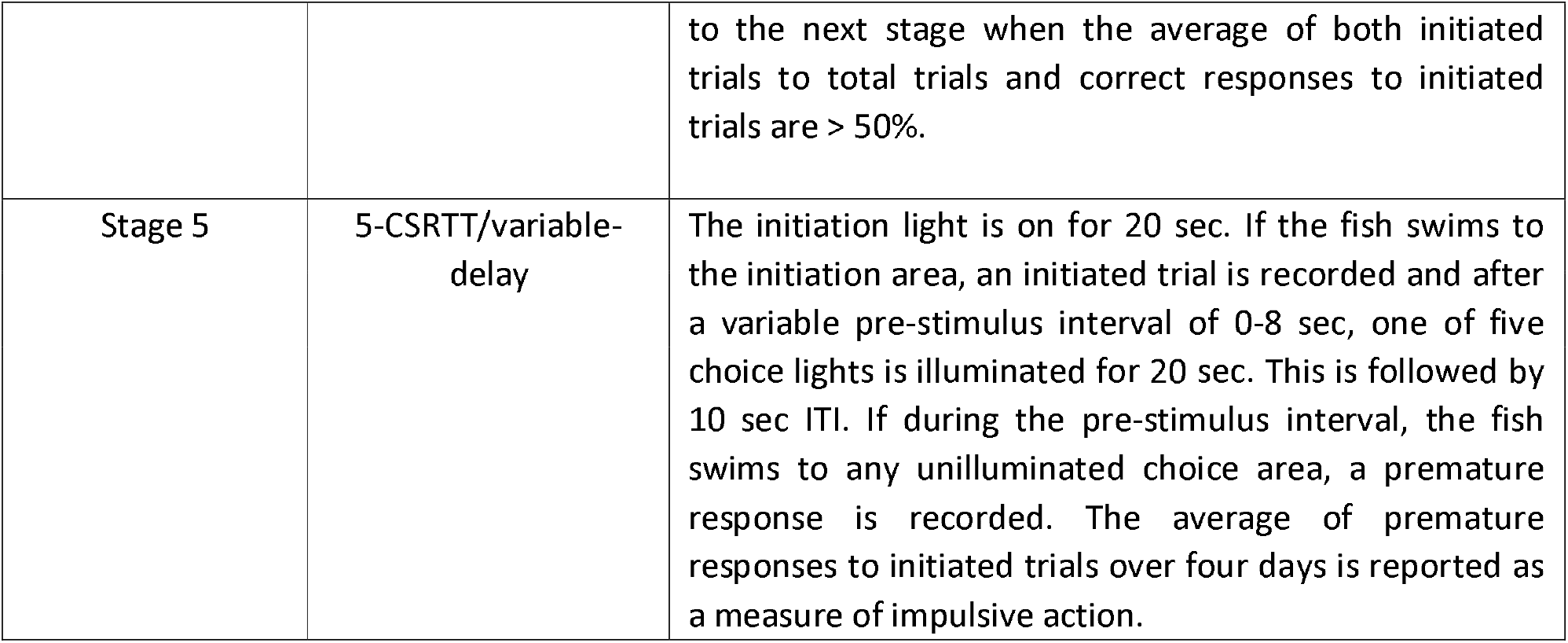
5-Choice Serial Reaction Time Task stages (5-CSRTT) and corresponding descriptions. Source: https://zantiks.com

## Notes

https://github.com/AdeleLeg

### Competing Interest Statement

The authors have declared no competing interest.

### Summary of Updates

1) Title has changed 2) New experiments have been added (cortisol measurement, more HPI axis qPCRs, behavioural experiments in presence of TRKB agonist and antagonist 3) Methods and Discussion have been implemented according to the new experiments and results

